# B7-H3 as a Dual Clinically Relevant Checkpoint and Antibody Drug Conjugate Target Expressed Across Adenocarcinoma and Neuroendocrine Prostate Cancers

**DOI:** 10.1101/2025.01.21.632253

**Authors:** Shivang Sharma, Nikita Mundhara, Emirhan Tekoglu, Adrianna Amaral, Tamara L. Lotan, Pan Gu, Jun Luo, Ezra G. Baraban, Nathan A. Lack, Eugene Shenderov

## Abstract

**Background and Objectives:** *CD276* (B7-H3) has recently emerged as a promising presumptive immune checkpoint inhibitor (ICI) and a potential antibody-drug conjugate (ADC) target for prostate cancer (PCa). We evaluated B7-H3 and 15 other clinically relevant ADC and ICI targets for expression at the RNA and protein level across the PCa continuum—hormone-sensitive, castration-resistant, and neuroendocrine.

**Methods:** CCLE data analysis and western blot experiments were performed for quantifying RNA and protein expression variability across PCa cell lines. Inter- and intratumoral heterogeneity was evaluated by integrating single-cell RNA sequencing data across 595k cells and 102 patients, spanning the disease continuum. AR and B7-H3 knockouts of PCa cell lines were developed and investigated using R1881 and Enzalutamide to elucidate the AR-CD276 signaling pathway through qRT-PCR and flow cytometry.

**Key Findings:** B7-H3 showed high expression in tumor and myeloid cells (tumor microenvironment – TME), lowest heterogeneity across all ADC and ICI targets, and was negatively regulated by androgen signaling.

**Limitations:** Further validation of ADC and ICI target protein expression in human samples and more exhaustive exploration of the AR-CD276 signaling cascade is required.

**Conclusion:** B7-H3 demonstrates the least susceptibility to selective pressure due to its stable expression across PCa disease states.

**Clinical Implications:** B7-H3 represents an important ADC target for PCa due to its potential to minimize drug resistance and likely ability to be a valid target across the prostate cancer continuum. Additionally, its expression in myeloid cells supports a dual role as an ICI, further enhancing its therapeutic relevance.

**Graphical Abstract:** **Discovery of optimal checkpoints and antibody drug conjugates (ADCs) in Prostate Cancer (PCa).**

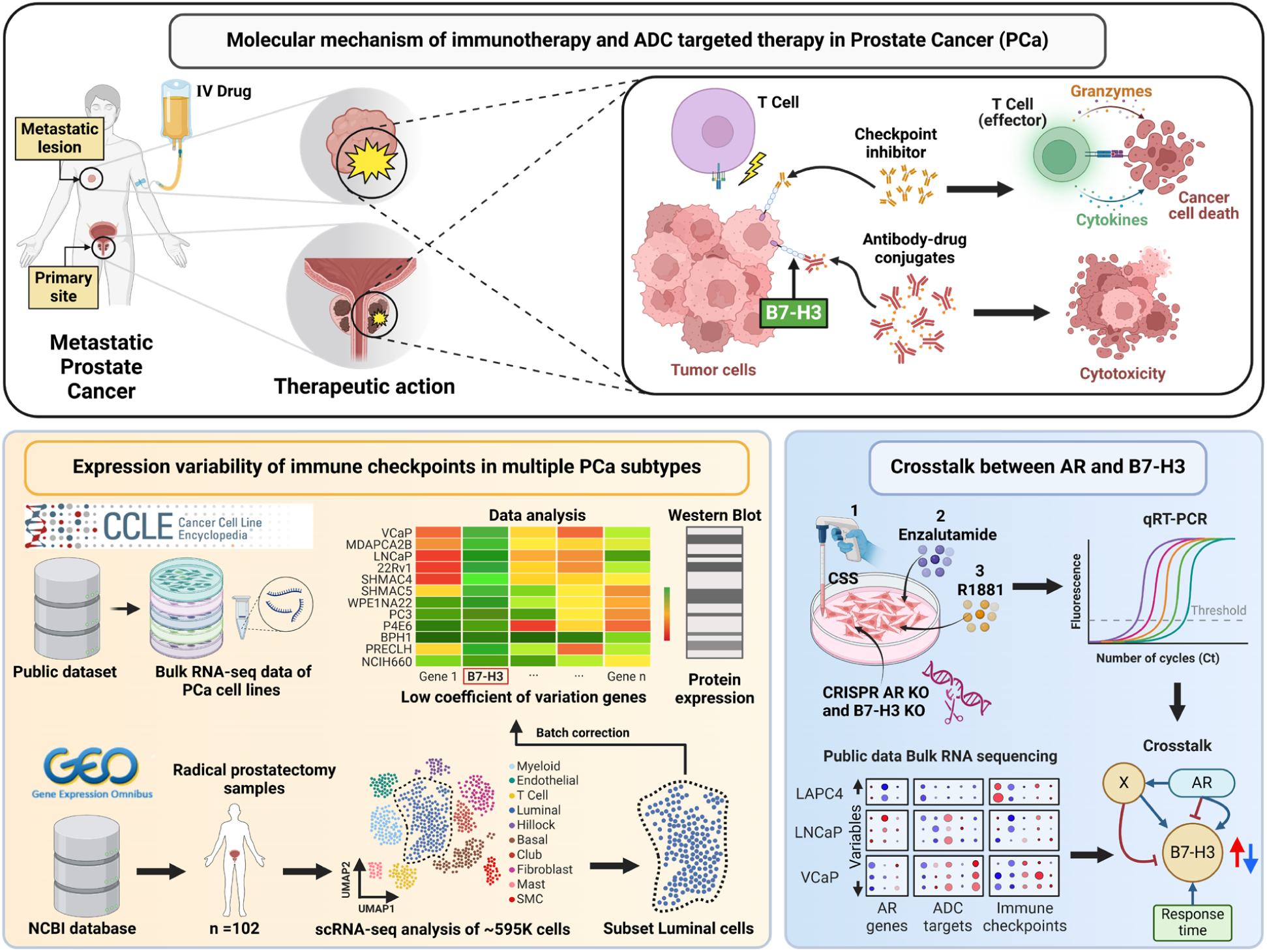

**Patient Summary:** In this study, we demonstrate that prostate cancer expresses high amounts of a cell surface protein called B7-H3 both when first diagnosed and throughout various stages of metastatic disease. Furthermore, we demonstrate a clear and yet to be fully resolved cross-talk between B7-H3 and the androgen signaling pathway central to prostate cancer development. We conclude that B7-H3 is therefore a highly promising therapeutic cell surface protein expressed by prostate cancer due to its stable expression potentially leading to low selective pressure and reduced drug resistance—thereby having significant implications for clinical trial design and patient inclusion considerations.

**Advancing Practice:** PCa is a highly heterogeneous disease with an increasingly wide treatment landscape, governed by various factors including clinical covariates, histopathological features, and molecular factors. B7-H3 has recently emerged as a promising clinical target for immunotherapy in the localized setting and antibody drug targeting in the metastatic setting. Despite the clinical significance, B7-H3 expression has not yet been thoroughly evaluated against other clinically relevant immune checkpoint and antibody drug targets, especially along the continuum of PCa treatment stages—hormone sensitive, castration resistant, and neuroendocrine. To our knowledge, this is the first study to characterize B7-H3 in this continuum using representative cell lines and the largest collection of PCa patient single cell sequencing data yet assembled in the literature, thus accounting for the substantial heterogeneity across PCa disease states as well as between datasets.

**Take Home Message:** This article explores the therapeutic relevance of clinical stage ADC targets and immune checkpoints for Prostate Cancer (PCa). We show that B7-H3 is a highly promising therapeutic candidate for PCa due to its stable expression across various disease states.

## INTRODUCTION

Prostate cancer (PCa) is the most commonly occurring non-skin cancer in men world-wide with one in eight men in the United States predicted to develop PCa, and continues to be the second most common cause of cancer-related death [1, 2]. Advanced prostate cancer is characterized by low immune infiltration, plasticity of tumor cells, and molecular heterogeneity presenting barriers to effective treatment, rendering the disease incurable and urgently requiring new treatment modalities[3]. Recently, Lutetium-177-PSMA-617 radiotheranostic was approved for metastatic castration resistant prostate cancer, but PSMA expression is required and limits efficacy to the AR-positive disease setting[4]. Unfortunately, immunotherapy has had limited success without PCa-specific approvals of immune checkpoint inhibitor (ICI) therapies. New immune checkpoint targets with increased activity in PCa and surface targets for chemotherapy delivery through antibody-drug conjugates are urgently needed across the PCa continuum of advanced disease, including neuroendocrine variant PCa (NEPC) whether de novo or treatment induced[5].

Several checkpoints have been identified as crucial to immune activation/suppression including: PD-1, PD-L1, PD-L2, lymphocyte activation gene 3 (LAG-3), T cell immunoreceptor with Ig and ITIM domains (TIGIT), tumor necrosis factor receptor superfamily 4 (OX40), 4-1BB (*CD137*) and cytotoxic T-lymphocyte associated protein 4 (CTLA4). Of these, three checkpoints now have FDA approval including PD-1/PD-L1, CTLA-4, and most recently LAG-3[6]. Crucially, none of these have had PCa-specific FDA approvals and there have been multiple negative single agent phase 3 trials and phase 2 combination trials [7–9]. For ADC targets in PCa, there are several tumor-expressed targets that have shown clinical promise including six-transmembrane epithelial antigen of prostate-1 and 2 (STEAP-1 and 2), prostate-specific membrane antigen (PSMA), nectin cell adhesion molecule 1 (NECTIN-1), trophoblast cell surface antigen-2 (TROP-2), and delta-like protein 3 (DLL-3)[10]. Each has been positioned in clinical development for different androgen-sensitive versus castration-resistant versus NEPC disease states, but the role for each so far has been limited due to tumor heterogeneity and what appears to be treatment-specific selective pressure to select for lower expressing subclones [11, 12].

B7 homolog 3 (B7-H3; *CD276*) is a presumptive immune checkpoint [13] whose binding partner remains to be elucidated. While many checkpoint proteins like PD-L1 are expressed at low levels in PCa, B7-H3 is highly expressed and associated with a worse prognosis[14]. We have recently shown that monotherapy in the neoadjuvant PCa setting with an Fc-engineered B7-H3 targeted antibody that mediates antibody-dependent cellular cytotoxicity was safe, induced immune activation, and potentially has clinical activity[15]. Furthermore, we and others have described high B7-H3 expression in advanced PCa as well as having an as-of-yet poorly characterized relationship to the androgen receptor (AR)[16, 17].

The objectives of this study were to explore relevant ADC and immune checkpoints in distinct PCa subtypes ranging from adenocarcinoma to NEPC to characterize their clinical heterogeneity and identify their possible relationship with AR. We have compiled ADC targets that are under early-to-late-stage clinical development for PCa and have explored the mRNA and protein expression of these targets in AR-independent/dependent cell lines, with and without treatment with AR activators/inhibitors. We have also investigated publicly available single-cell RNA sequencing data (comprising ∼595K cells, 102 patients and 124 samples) for healthy prostate tissue, localized PCa and adjacent benign prostatic tissue, and metastatic tumors for these ADC targets. The expression of selected targets of interest were validated using RT-PCR and western blot for AR independent/dependent cell lines. From these large datasets, we demonstrated that B7-H3 is uniformly highly expressed across all PCa subtypes, with the lowest coefficient of variation across current ADC and ICI targets. Furthermore, B7-H3 appears to be a downstream target of AR regulation through an indirect mechanism. Taken together these findings indicate that B7-H3 might serve as a therapeutic target across the PCa continuum and allow both ICI and ADC strategies to be pursued both as single agent and combination therapies.

## MATERIALS AND METHODS

### Cell culture

LNCaP, DU145, VCaP, 22RV1, PC3, NCI-H660, and PrECLH cell lines were authenticated for mycoplasma and STR analysis. LNCaP B7-H3 Scramble and LNCaP B7-H3 KO cells were created using abm’s All-in-one sgRNA-Cas9-puromycin vector. They were cultured in RPMI media 10% FBS. LNCaP95 is a strain of LNCaP cultured in RPMI media supplemented with 10% charcoal-stripped (androgen-depleted) FBS for several generations. LNCaP95, LNCaP95 AR Scramble, and LNCaP95 AR KO cells used in this study were created in Dr. Jun Luo’s lab [18]. The cultures were grown in a 37°C humidified incubator with 5% CO_2_. All reagents, antibodies, and TaqMan probes their company and Catalog numbers are provided in **Table S1**.

### Western blotting

Protein lysates from cell pellets were used to quantify the expression of B7-H3, AR and PSMA from various PCa cell lines. H69 (Lung cancer) was used as cell line control. LNCaP WT, LNCaP B7-H3 Scramble, LNCaP B7-H3 KO, LNCaP 95, LNCaP95 AR Scramble, and LNCaP95 AR KO cell pellets were confirmed for their respective KOs. LNCaP, VCaP, 22RV1, and LAPC4 cells were treated with R1881 (10 nM) for 96 h. Protein lysates were prepared using cell lysis buffer supplemented with protease and phosphatase inhibitor. Western blotting was performed the probed for B7-H3, PSMA, AR. β-actin was used as a loading control. The chemiluminescent along with fluorescent signals were measured using a Bio-Rad chemidoc imaging system and analyzed by ImageJ software. Date represents the mean of 3 independent experiments.

### Real-time PCR

LNCaP and DU145 cells were treated with media containing charcoal-stripped FBS (CSS), CSS + R1881 (10 nM) and Enzalutamide (10 μM) for 96h. RNA isolation was performed using a Qiagen RNeasy kit, followed by cDNA preparation using a High-Capacity cDNA Reverse Transcription Kit with RNase Inhibitor according to the manufacturer’s protocol. TaqMan assay probes c-MET, CD276, AR, KLK2, and GAPDH were used to quantify the mRNA expression on Applied Biosystems QuantStudio 3. Data normalized to GAPDH and represented relative to WT or no treatment control. Date represents the mean of 3 independent experiments.

### Flow cytometry

Cells were detached using a cell stripper, blocked by Fc blocker, and surface staining for B7-H3 using Enoblituzumab AF647 was performed. Dead cells were excluded from the analysis using the LIVE/DEAD™ Fixable Aqua Dead Cell stain kit. For intracellular staining of AR AF488, cells were fixed and permeabilized using an FOXP3 transcription factor staining kit. Samples were measured on a 5-laser Aurora spectral flow cytometer (Cytek Biosciences, Fremont, CA, USA) and analyzed using FCS Express Software. An example of flow gating strategy is provided in **Figure S1.** Date represents the mean of 3 independent experiments.

### Bulk RNA-seq data analysis

Raw data was obtained from GEO (Gene Expression Omnibus) and described in **Table S2**. Baseline expression data of all protein coding genes reported as log2(TPM+1) were obtained from the CCLE/DepMap portal (23Q2 release)[19] and filtered to include only the prostatic cell lines. Expression values in the heatmaps were obtained directly from the filtered table while AR and NE scores [20, 21] for each cell line were calculated with GSVA (default parameters)[22].

Differential expression analysis for each sample was performed using an in-house pipeline. Reads mapping to the human genome and gene quantification was performed using primary assembly of GRCh38.p13 in ENSEMBL release 109 and STAR (2.7.10b)[23]. Differential expression analysis compared to control samples was performed using DESeq2[24] and log2 fold change values shrunk using the ‘ashr’ algorithm were used for plotting.

### scRNA-seq data analysis

scRNA-seq datasets, described in **Table S3**, were downloaded from GEO. Seurat package[25] and R programming language was used for the entire analysis. First, the datasets were individually analyzed to remove cells with gene counts lower than 500 and mitochondrial content > 25%. The filtered cells were then normalized with SCTransform and integrated at dataset level using Harmony algorithm using top 4000 variable genes. The integrated data was clustered, using the first 40 principal components, resolution of 4 and Leiden clustering algorithm, onto a UMAP. Doublets were filtered from the integrated dataset using the DoubletFinder[26] package and a multiplet rate of 8%. The final dataset comprised of ∼595K cells. The clusters were annotated for characteristic prostate cell types using ensemble canonical marker genes mentioned in **Table S4**. Normalized, log-transformed expression of Expression of ADC and ICI targets converted to densities using kernel density estimation before plotting on plotted on UMAP to capture overall expression of cells at a given (x, y) coordinate pair in the UMAP.

The integrated, annotated dataset was subsetted for high AR and NE score cell types (Luminal and Neuroendocrine) and converted into pseudo-bulk data. Some patients contributed to more than one sample through tumor and matched normal, amounting to 124 samples from 102 patients. The pseudo-bulk dataset was log-normalized for size factors and batch effects were regressed out across datasets using limma package, while preserving biological variability by providing sample-level subtype information as experimental conditions to be preserved. The batch-corrected data was scored for AR and NE genesets and used to analyze the log-expression values of the relevant ADC and ICI targets. Intratumoral heterogeneity was calculated on Luminal and Neuroendocrine cell populations for each sample in the single-cell dataset. It was calculated as the percent coefficient of variation of the target’s normalized expression across all the cells in that given sample. Log-normalized B7-H3 expression between Luminal and Basal cells were plotted as box plot and Welch t-test was performed to evaluate statistical significance.

### B7-H3 Immunohistochemistry

The B7-H3 protein expression was examined in a cohort of 22 biopsy or transurethral prostate resection specimens from patients with small cell neuroendocrine prostate carcinoma. Seven patients had concomitant conventional adenocarcinoma represented within the specimen. Three to twelve 1 mm tissue cores from each tumor were punched to create a TMA block [27, 28]. We stained the TMA slide for B7-H3 using a previously validated protocol on the Ventana Benchmark (Ventana/Roche, Tucson, AZ) [17]. The tumor areas were annotated and B7-H3 staining was scored using Membrane v1.7 algorithm in HALO® (IndicaLabs).

## RESULTS

### B7-H3 is more uniformly expressed among clinically relevant ADC and ICI targets in AR independent and dependent cell lines

We investigated ADC and immune checkpoint targets in active clinical development for all major subtypes of PCa. Baseline RNA expression from the Cancer Cell Line Encyclopedia (CCLE) was examined for AR-dependent/independent, primary, metastatic, and NEPC, as well as normal and benign prostate cell lines. The cell lines were scored for neuroendocrine (NE) and androgen-receptor pathway (AR) signatures using published gene sets (Methods). Similar to published work, AR, PSMA, STEAP1 and STEAP2 are highly expressed in VCaP, LNCaP, 22Rv1, and MDA-PCa-2b, and correlate well with the AR pathway activity scores whereas they have lower expression in AR-negative cell line models (**Figure 1A**). Conversely, models representing prostatic small cell or neuroendocrine carcinoma such as NCI-H660 express a relatively low or lack entirely these ADC targets and instead express other genes such as *DLL3* [29]. Similarly, **Figure 1B** shows the expression of clinically relevant immune checkpoint molecules. B7-H3 expression has the lowest coefficient of variation (CV) amongst all the ADC and the ICI target candidates shown in **Figure 1A and 1B** and described in **Table S5**. Immune checkpoints like TIGIT, OX40, 4-1BB and CTLA4, which are expressed primarily in immune cells, were utilized primarily as negative/low expression controls. These observations were orthogonally validated by comparing the protein expression (western blot) of PSMA, AR, B7-H3, PD-L1, PD-L2, STEAP1, STEAP2, TROP2, NECTIN1, PSCA, LAG3, DLL3, and 41BB across AR-positive adenocarcinomas (LNCaP, VCaP, 22Rv1), AR-negative adenocarcinomas (PC3, DU145), a prostatic neuroendocrine carcinoma (NCI-H660), and a lung carcinoma (negative control; NCI-H69 (**Figure 2**). The RNA and protein expression data were mostly correlated. Overall, we observed that B7-H3 protein was expressed in all cell lines. AR-positive LNCaP, VCaP, and 22RV1 cell lines had the greatest expression, AR-negative DU145 and PC3 cell lines demonstrated lower expression, and the neuroendocrine H660 cell line somewhat in-between. PSMA expression was limited to AR-positive prostatic adenocarcinomas.

**Figure 1.**
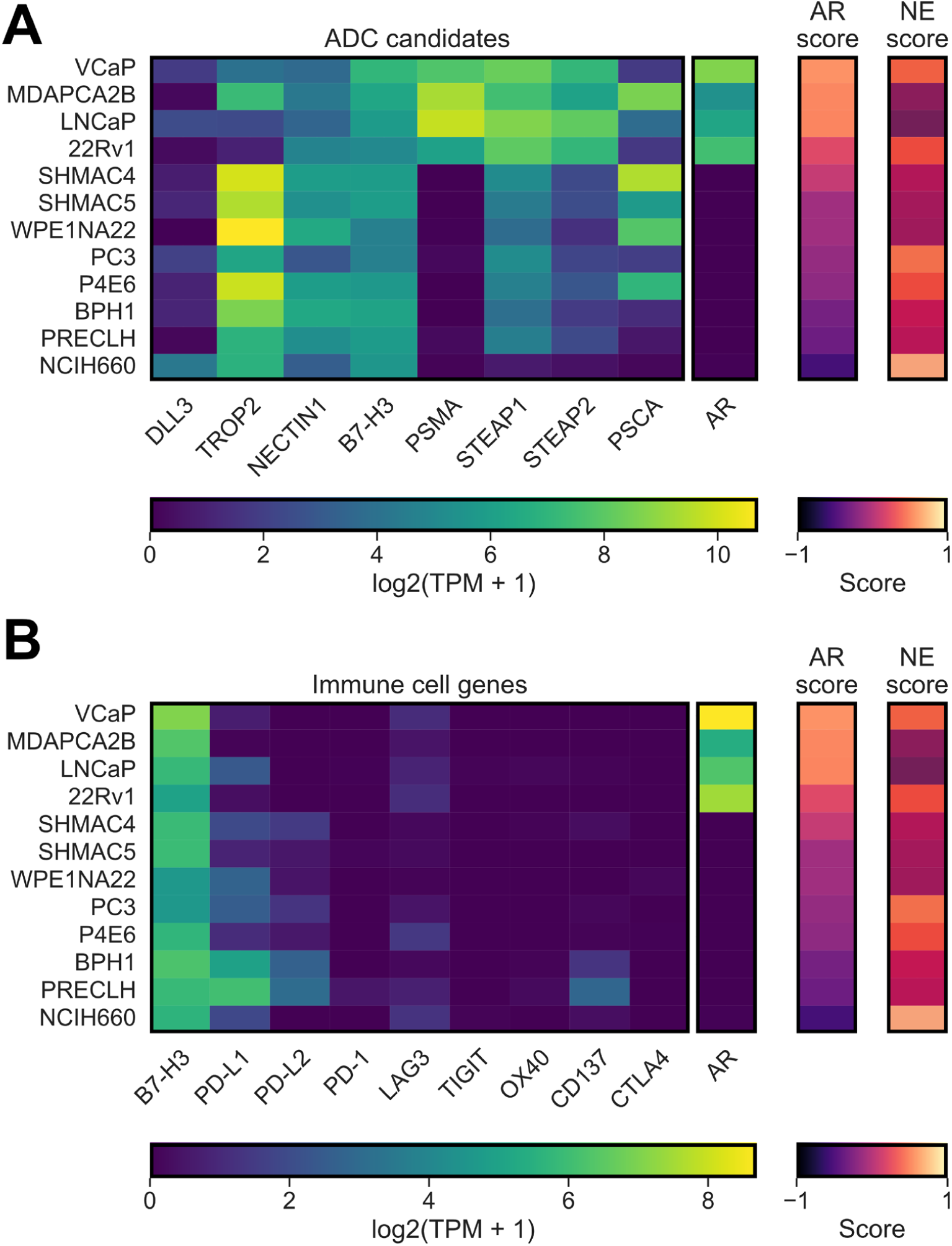
RNA expression of selected ADC and immune targets in active clinical development. Bulk RNA sequencing data obtained from CCLE for various cell lines (y-axis), describing log(TPM+1) of **A)** important ADC targets and **B)** important immune checkpoints. The AR and NE scores were calculated using the log TPM, with higher score (Methods) indicating higher expression of AR or NE associated genes in the corresponding cell lines. B7-H3 is included as both an ADC target candidate and immune cell gene.

**Figure 2.**
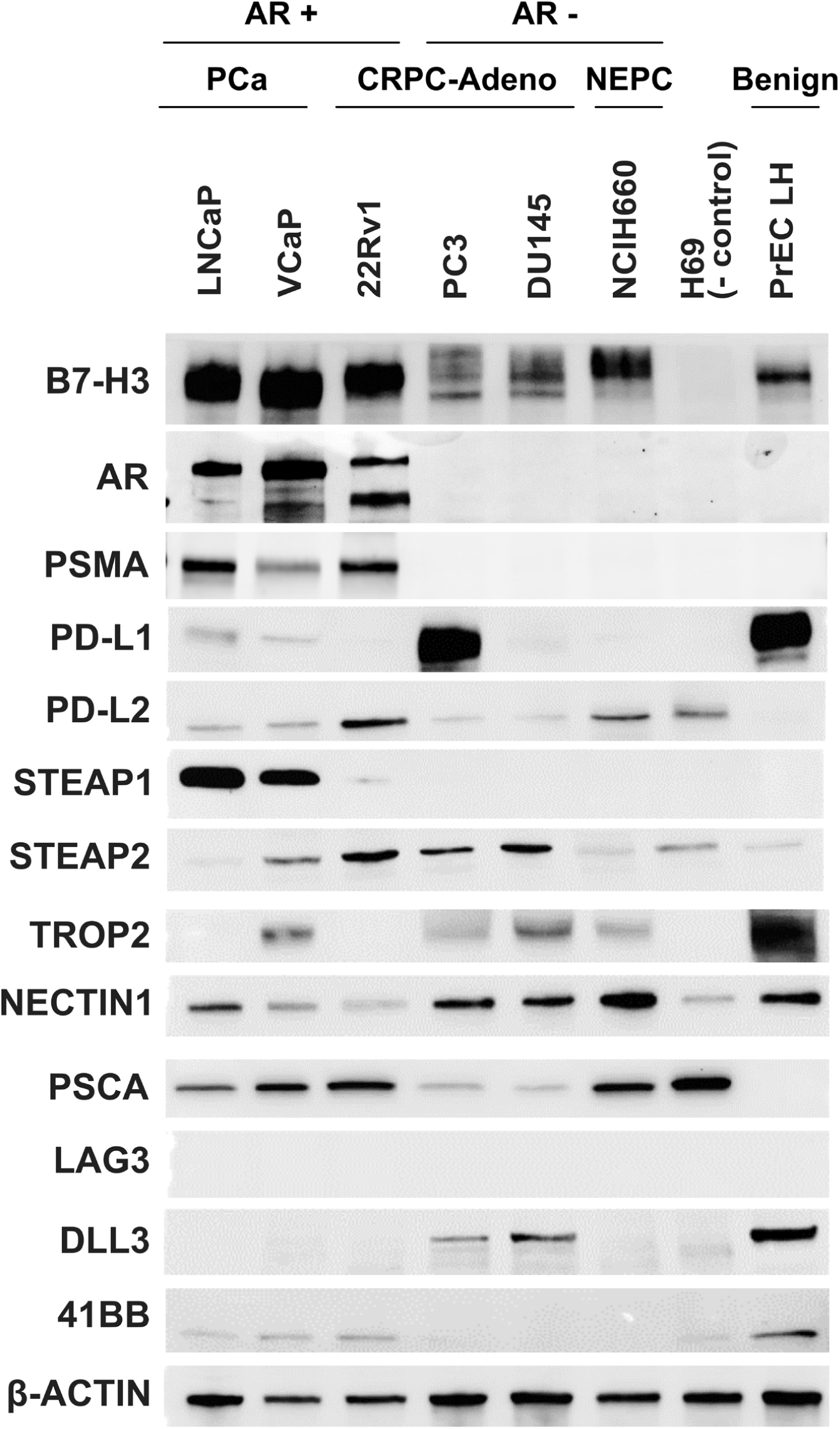
Protein expression of selected ADC and immune targets in active clinical development. Western blot depicting the protein expression of selected ADC and immune targets in LNCaP, VCaP, 22Rv1, PC3, DU145, NCI-H660 prostate cancer, and PrEC LH prostate non-cancerous cell lines. H69 (lung carcinoma) is a negative control for B7-H3 expression. β-actin was used as a loading control (N=3).

### B7-H3 exhibits highly stable RNA expression across multiple pathophysiological subtypes of PCa and multiple patients

RNA expression of clinically relevant ADC and ICI targets in cell lines was validated by analyzing eleven high-quality scRNA-seq datasets of prostate tissue spanning across multiple pathological states, as described in **Table S3**. The quality-controlled (**Figure S2**), integrated and annotated dataset was used to visualize the expression of important ADC and ICI targets across different cell types, shown in **Figure 3A**, with the expression densities of luminal markers shown in **Figure S3**. We observed that B7-H3 (CD276), KLK3 and PSMA (FOLH1) are predominantly expressed in luminal cells while TROP-2 (TACSTD2) and NECTIN1 are expressed primarily in basal and club cells. Immune-related genes PD-1, PD-L1, PD-L2, LAG3 and OX40, are primarily expressed in the immune cell clusters (myeloid, mast or T cells), except for HLAs which are expressed on multiple cell types (**Figure S4**). HLAs also show a heterogeneous expression (with varying degrees across different HLAs) in cell lines as shown in **Figure S5**. B7-H3 was the only examined protein abundantly expressed on both tumor and immune (myeloid) compartments.

**Figure 3.**
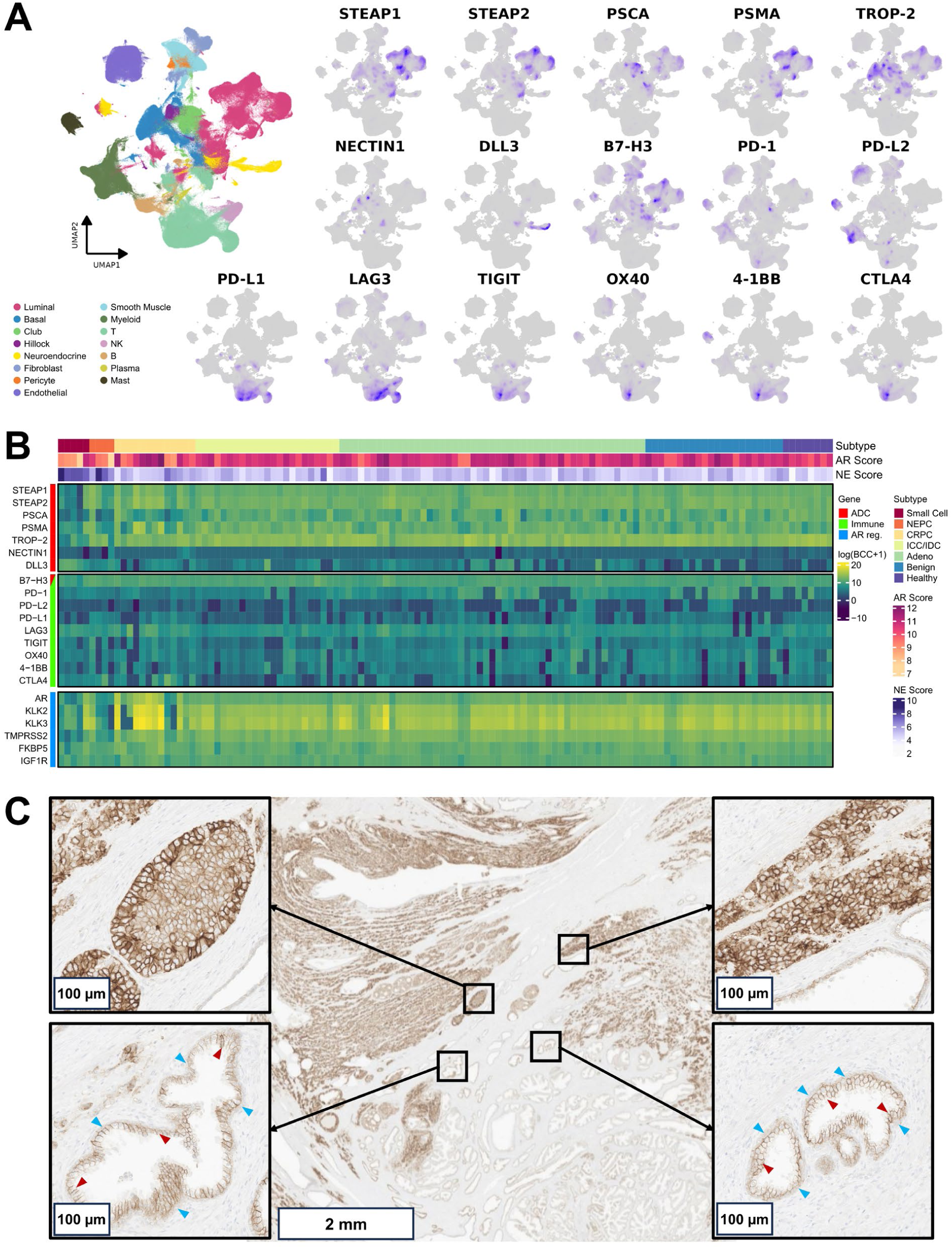
Publicly available scRNA-seq datasets of Healthy, Benign (matched normal), Adenocarcinoma (Adeno), Cribriform (ICC/IDC), Castration Resistant Prostate Cancer (CRPC), Neuroendocrine Prostate Cancer (NEPC) and Small Cell Prostate Cancer were integrated into a single dataset, comprising 102 patients, 124 samples and ∼595K cells, followed by clustering and annotation. (**A**) UMAP plot of integrated dataset annotated with major cell types found in prostate tissue. Log expression of important ADC targets and immune checkpoints are shown over individual UMAP density plots with purple color representing higher expression; (**B**) Integrated single cell dataset is subsetted for high AR and NE scoring cells and converted into pseudo-bulk data. Batch corrected log pseudo-bulk counts, log(BCC + 1), of ADC targets (red), immune checkpoints (green) and AR associated genes (blue) in the pseudo-bulk data are shown in heatmap, with each column representing a separate patient. The ‘Subtype’ bar on top clusters the patients into tumor subtypes, and AR and NE scores are calculated for each patient; (**C**) B7-H3 staining of representative primary PCa, showing benign and tumor regions. Prostatic adenocarcinoma (upper left and right inset) shows higher B7-H3 expression compared to adjacent benign prostatic glands (lower left and right inset). Within benign prostatic glands, higher expression of B7-H3 is evident in secretory cells adjacent to the glandular lumen (red arrowheads) compared to the minimal B7-H3 labeling observed in the underlying layer of basal cells (blue arrowheads). Samples marked as CRPC histological features of adenocarcinoma and no neuroendocrine or small-cell like features are observed.

Next, the integrated dataset was subsetted for cell types with high scores for androgen (AR) and neuroendocrine (NE) signatures. Epithelial subtypes and Neuroendocrine scored highest for AR and NE signatures (**Figure S6**) and were used to understand the variability in expression of important ADC targets across different subtypes of PCa. The subsetted single-cell dataset was transformed into pseudo-bulk data and batch-corrected (**Figure S7)** to compare the gene-expression profiles across multiple patients and PCa subtypes. Genes corresponding to important ADC targets were visualized for their variation across the disease types, as shown in **Figure 3B**. Consistent with analyzed cell lines B7-H3 showed the least amount of variation (CV = 6.23%), among the tumor-expressed markers, followed by IGF1R (CV = 6.78%) and FK1BP5 (CV (9.03%). In contrast, STEAP1, FKBP5 and TMPRSS2, all showed higher variation (**Table S5**). Among the immune-related genes, the variability in all the genes, except LAG3 (CV = 21.26%), exceeded 25% (**Table S5**). B7-H3 also showed low intratumoral heterogeneity across all the patients in the single-cell dataset, shown in **Figure S8**.

Expression of B7-H3 was evaluated across the entire integrated dataset containing all cell types, with emphasis on the tumor subtypes. Luminal cells demonstrated significantly higher expression of B7-H3 compared to basal cells (**Figure S9**). This was confirmed at the protein level by B7-H3 IHC performed on primary prostate tumors, which demonstrated differential B7-H3 expression between benign glands and tumor epithelial cells. Benign prostate glands demonstrated weak to moderate B7-H3 expression in secretory cells with minimal expression observed in underlying basal cells. In contrast, neoplastic glands showed diffuse and strong B7-H3 expression (**Figure 3C**). We examined a previously described set of neuroendocrine prostate carcinomas (NEPC) [27, 28]—frequently AR negative or AR independent—sampled with or without concurrent adenocarcinomas (**Figure S10A**). Comparing all the NEPC to the concurrent adenocarcinomas, there was a trend towards higher B7-H3 protein expression among the adenocarcinomas (**Figure S10B**). In a paired analysis, comparing concurrent adenocarcinoma and neuroendocrine components within an individual patient, a similar trend was observed (**Figure S10C**), but overall B7-H3 protein expression was relatively stable.

### Inhibiting AR increases B7-H3 expression whereas B7-H3 KO causes no change in AR expression in LNCaP cells

AR is a ligand-induced transcription factor that is activated by androgens like dihydrotestosterone and translocates to the nucleus where it binds to genomic DNA and regulates the expression of a variety of genes. There is conflicting evidence about the role of AR in B7H3 expression [14, 17]. In this work, we observed that B7-H3 is ubiquitously expressed in all PCa cell lines and patients (**Figures 2 and 3B**). To understand the effect of AR activation on B7-H3 expression, we analyzed multiple bulk RNA-seq datasets of LNCaP, LAPC4 and VCaP with an acute androgen treatment, treatment durations (4 – 24 h) and dosages (**Figure 4A**). As expected, the AR induced genes were acutely upregulated in all three models while AR was downregulated, especially in VCaP owing to the high amount of AR present and strong negative feedback loop [30]. None of the ICI targets investigated were differentially expressed across all three models following androgen induction under any condition. While some ADC targets were differentially expressed, there was a large variance and disagreements across models and samples. B7-H3 expression, on the other hand, did not change consistently across models, with the exception of a slightly more significant downregulation in VCaP, presumably due to the several fold difference in AR expression relative to the other two models.

**Figure 4.**
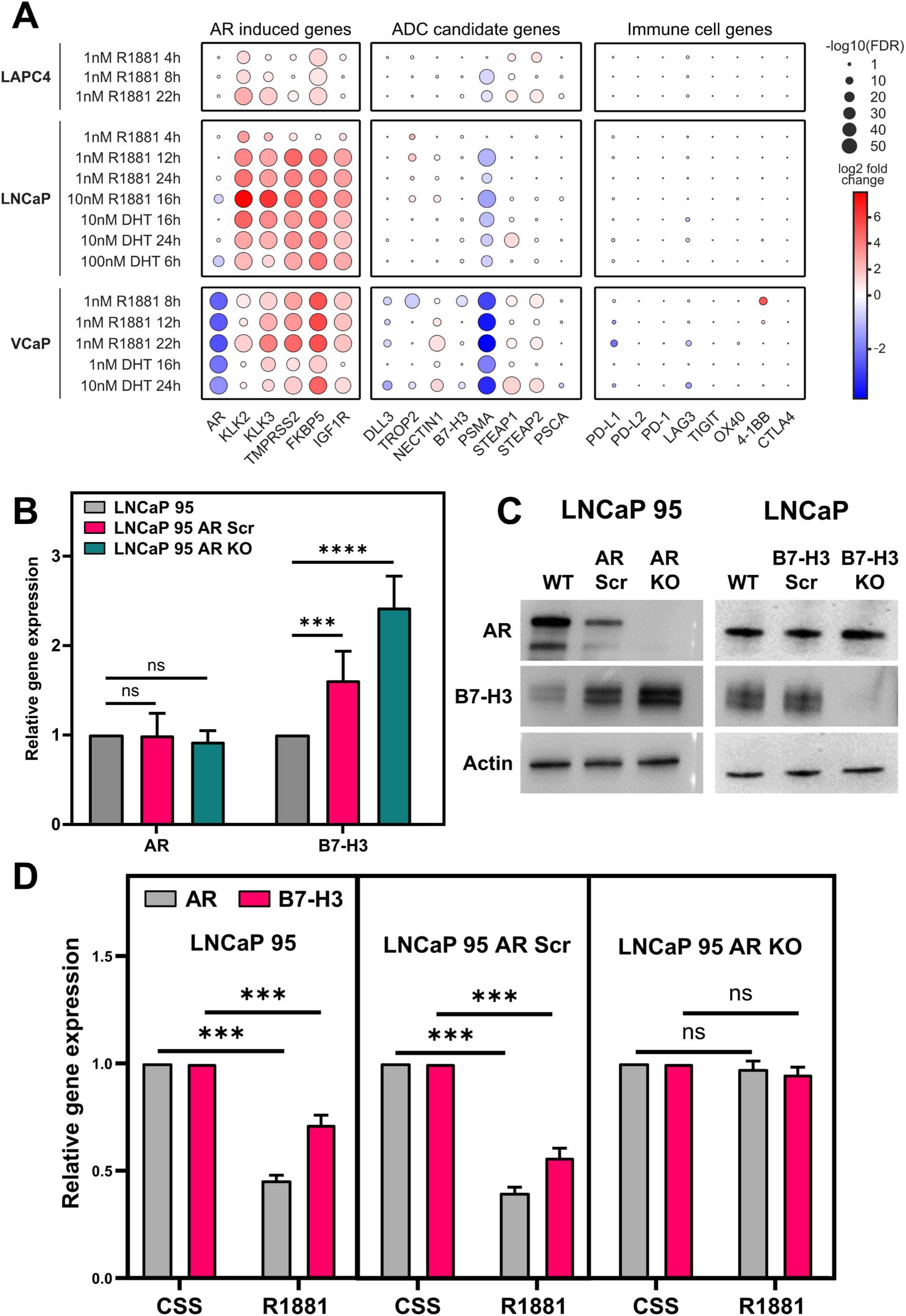
(**A**) Differential gene expression of AR regulated genes, ADC candidate genes and immune cell genes of interest with androgen treatment in prostate cancer cell lines. Fold-change versus control is shown in expression of AR regulated genes, immune checkpoints and ADC candidates in LAPC4, LNCaP and VCaP cell lines upon treatment with different concentrations and treatment durations of R1881 and DHT. (**B**) Relative gene expression of AR and B7-H3 in LNCaP 95, LNCaP 95 AR Scr, and LNCaP 95 AR KO cell lines. All the data points are normalized to GAPDH control (N=3). (**C**) Representative blots depicting the AR and B7-H3 protein expression in LNCaP 95, LNCaP 95 AR Scr, LNCaP 95 AR KO, LNCaP WT, LNCaP B7-H3 Scr, and LNCaP B7-H3 KO cell lines (N=3). Actin was used as a loading control. (**D**) Relative gene expression of AR and B7-H3 in LNCaP 95, LNCaP 95 AR Scr, and LNCaP 95 AR KO cell lines in 10 nM R1881 treatment for 96 h with respect to CSS untreated controls. All the data points are normalized to GAPDH control (N=3). The error bars represent SD (ns = non-significant, **** = p value < 0.0001, *** = p value < 0.001).

To further probe the relationship between AR and B7-H3, we utilized stable CRISPR KOs of AR and B7-H3 in this study. AR KO LNCaP95 cells displayed no AR expression at the protein level, however AR was detected at the RNA level. B7-H3 expression was significantly increased in LNCaP95 AR KO cell lines compared to LNCaP95 and LNCaP95 AR scramble at both RNA (**Figure 4B**) and protein levels (**Figure 4C**). However, LNCaP B7-H3 KOs did not show any discernible change in AR expression compared to LNCaP and LNCaP B7-H3 scramble cells (**Figure 4C**), quantification represented in **Figures S11A & S11B**. This data suggests that B7-H3 is a downstream target of AR with further research still required to elucidate the mechanism.

### External androgen, androgen deprivation and AR antagonist alter B7-H3 expression in only AR+ PCa cell lines

To understand AR mediated B7-H3 regulation, LNCaP95, LNCaP95 AR scramble, and LNCaP95 AR KO were treated with R1881. R1881 is a synthetic androgen that stimulates AR signaling. Amongst the primary downstream targets of AR, Kallikrein 2 (KLK2) is a positively regulated gene, whereas c-Met is negatively regulated [31] and these were used as controls (**Figure S11C**). R1881 treatment significantly suppresses B7-H3 expression in LNCaP95, and LNCaP95 AR scramble cells, but not in LNCaP95 AR KO cell lines (**Figure 4D**), suggesting that the negative regulation of B7-H3 is a consequence of AR activation.

Further, androgen deprivation by CSS treatment - that decreases AR signaling - increases B7-H3 expression in AR (+) LNCaP cells but not in AR (-) DU145 cells, providing another layer of support to the AR/B7-H3 regulation axis. Moreover, CSS supplemented with R1881 decreases B7-H3 expression with respect to only CSS-treated cells (**Figure 5A**). A similar trend was observed at the protein level in AR (+) LNCaP and 22Rv1 cells by flow cytometry (**Figure 5B**). Given the changes in B7-H3 expression were observed in only AR (+) cell lines, we confirmed that R1881 (10 nM) for 96 h reduces B7-H3 expression in four AR (+) PCa cell lines - LNCaP, LAPC4, 22Rv1 and VCaP (**Figure 5C**), quantification represented in **Figure 5D**.

**Figure 5.**
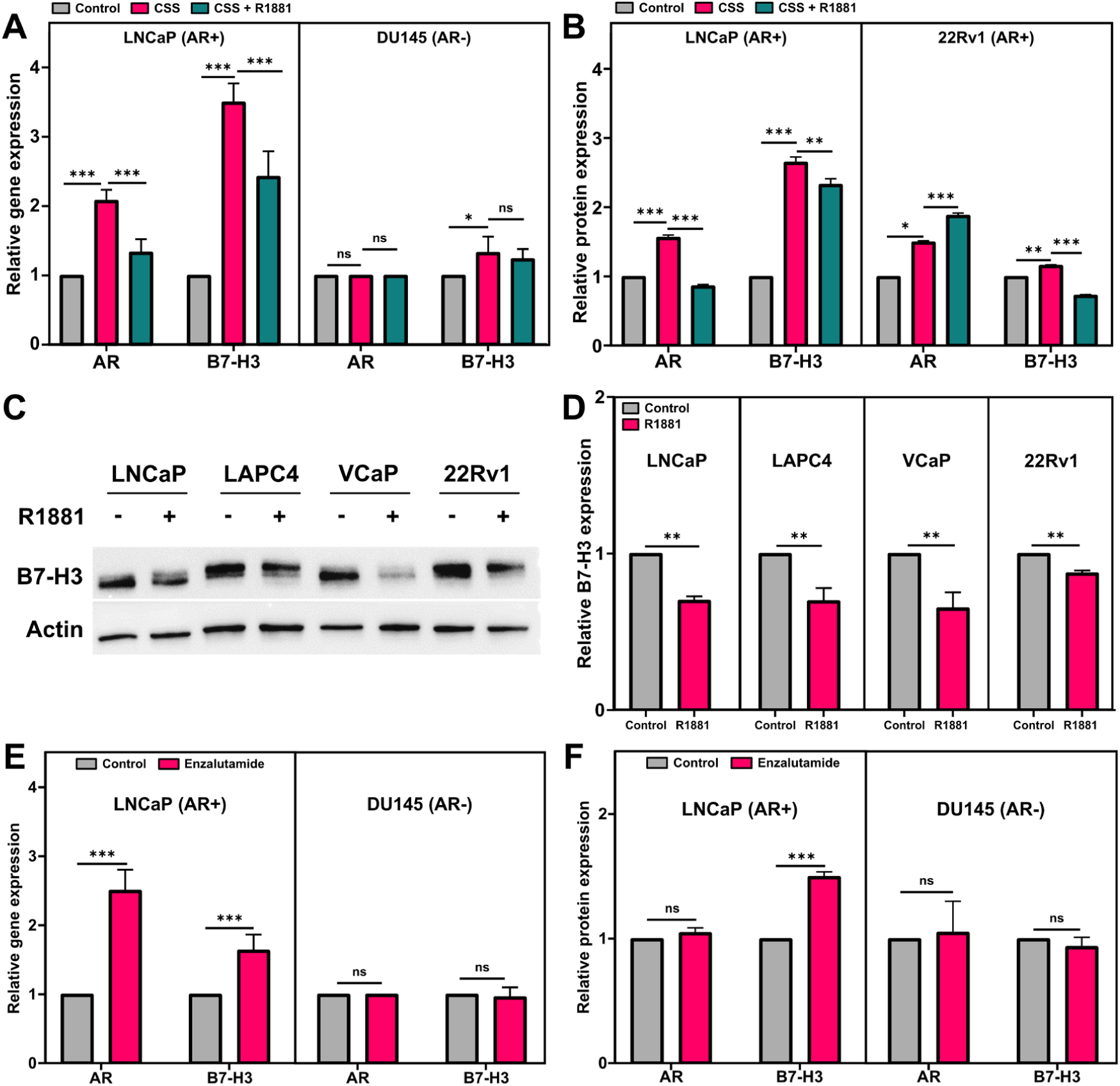
(**A**) Relative gene expression of AR, B7-H3 in LNCaP and DU145 cells. Bar graph representing the relative gene expression upon treatment with CSS, CSS + R1881 (10 nM) for 96 h (N=3) with respect to RPMI untreated controls. All the data points are normalized to GAPDH control. (**B**) Relative protein expression (by flow cytometry) of AR, B7-H3 in LNCaP and 22Rv1 cells upon treatment with CSS, CSS + R1881 (10 nM) for 96 h with respect to RPMI untreated controls (N=3). (**C**) Representative blots displaying B7-H3 expression upon R1881 (10 nM) treatment for 96 h in LNCaP, LAPC4, VCaP, and 22Rv1 cell lines (N=3). Actin was used as loading control (**D**) Densitometric quantification of the western blot shown in Figure 4c. The data is represented with respect to untreated controls. All data points are normalized to vinculin (**E**) Relative gene expression of AR, B7-H3 in LNCaP and DU145 cells. Bar graph representing the relative gene expression upon treatment with Enzalutamide (10 uM) for 96 h (N=3) with respect to RPMI untreated controls. All the data points are normalized to GAPDH control. (**F**) Relative protein expression (by flow cytometry) of AR, B7-H3 in LNCaP and 22RV1 cells upon treatment with Enzalutamide (10 uM) for 96 h with respect to RPMI untreated controls (N=3). The error bars represent SD (ns = non-significant, *** = p value < 0.001, ** = p value < 0.01, * = p value < 0.05).

To further confirm the observed inverse association between AR signaling and B7-H3, we administered enzalutamide treatment in LNCaP and DU145 cells. Enzalutamide is an anti-androgen utilized in the clinic, which sequesters the AR from translocating into the nucleus and thereby inhibits AR-regulated transcriptional activity. Enzalutamide treatment increased B7-H3 expression at both RNA (**Figure 5E**) and protein levels (**Figure 5F**) in AR (+) LNCaP cells. This independently endorses that AR negatively regulates B7-H3. There were no significant changes observed in AR (-) DU145 cells.

## DISCUSSION

Our work corroborates the earlier findings that the expression of PSMA, STEAP1, TROP2, and DLL3 fluctuates across the entire spectrum of PCa. While TROP2 is primarily expressed across CRPC molecular subtypes but absent in NEPC, STEAP1 exhibits a wider expression than PSMA in the context of lethal mCRPC despite not always being expressed at higher levels across all mCRPC tissues [32]. On the other hand, DLL3 is mainly expressed on NEPC [33]. B7-H3 was shown to have high expression both inside and across all metastatic prostate cancers in a prior work that focused on assessing gene expression using digital spatial profiling [34]. By removing any spatial biases and confirming that B7-H3 is the ADC and ICI target expressed with the least heterogeneity across the entire spectrum of PCa, our work has expanded on the concept using RNA and protein level data across the entire spectrum of prostate cancer from cell lines to single cell level patients.

Among all the investigated ADC and ICI targets, B7-H3 shows the most uniform RNA expression, across all the PCa cell lines and patient tumor samples. Similar trends are observed for the protein expression across PCa cell lines. We found that the coefficient of variation (CV) was lower in single-cell analysis compared to the cell line data. This could primarily be due to the higher number of samples in the integrated single-cell dataset (number of samples = 124) compared to the CCLE data (1 sample per cell line from 12 cell lines). This is in contrast to the highly variable protein expression of targets like PSMA and AR, which also show similar heterogeneity at the level of RNA expression, both in the cell lines and patient samples. Qualitative agreement between RNA and protein expression profiles enabled use of RNA expression as a surrogate for protein expression and indicates that most of the ADC targets investigated would likely exhibit heterogeneous protein expression, when analyzed across PCa subtypes. Intratumoral heterogeneity is one of the reasons for clonal evolution of PCa and other cancers [12], especially under internal or external selective pressures like hypoxia or therapeutic intervention. The uniform, strong expression of B7-H3—albeit with lower and higher levels between AR-negative and positive cell lines—noted herein make it an ideal candidate for a wider cohort of PCa patients and limit subclonal escape under selection pressure as B7-H3 null prostate cancer was not observed in our study.

Other common targets, such as PSMA and STEAP1, have been shown to be selectively down-regulated [35, 36], leading to therapeutic resistance. Some cancer cells in hormone-sensitive and insensitive prostate tumors have shown low expression of B7-H3 but these could potentially remain targetable by bystander activity as recently described [16]. Our findings have implications for inclusion/exclusion criteria for B7-H3 prostate cancer clinical trial enrollment as some trials actively exclude subjects exhibiting histopathological signatures of NEPC, and these patients should not be excluded.

The majority of PCa cases are androgen-dependent [37]. In these tumors the AR acutely induces the expression of hundreds of genes (direct androgen regulation). These differentially expressed target genes then go on to affect the expression of other genes (indirect androgen regulation). We and others have shown a plausible relationship between B7-H3 and AR [14, 17]. Mendes *et al*. suggested that androgen signaling may positively regulate B7-H3 expression in patient biopsies whereas Benzon *et al*. demonstrated that the presence of androgen dampens B7-H3 expression in LNCaP cells. Due to these conflicting observations, we aimed to explore the direct and indirect effects of androgen on B7-H3 expression using multiple cell lines, knockout models, and androgen agonist-antagonist treatment conditions. We observed that B7-H3 is likely indirectly regulated by androgen treatment in AR-positive tumors. Our results with LNCaP AR KOs and B7-H3 KOs highlighted that B7-H3 is a downstream target of AR while the B7-H3 does not impact AR expression (**Figure 4C**). Further, LNCaP WT, AR scrambles, and KOs confirm R1881 decreases B7-H3 expression as observed by Benzon *et al*. However, these changes were only observed in the presence of AR (**Figure 4D**). No change in B7-H3 expression was observed following acute treatment of androgen suggesting that this does not occur via direct AR regulation (**Figure 4A**). However, following longer treatment (96hrs) LNCaP, VCaP, 22Rv1, and LAPC4 (AR positive cell lines) displayed a decrease in B7-H3 expression, but DU145 (AR negative cells) showed no significant changes upon R1881 treatment. Similarly, longer treatment of Enzalutamide caused an increase in B7-H3 expression in AR positive LNCaP cells but not in AR negative DU145 cells. Our finding that B7-H3 is negatively regulated by androgen, suggests that there may be increased sensitivity to B7-H3 directed therapies during androgen deprivation therapies and indicates a route toward future combination therapies.

Limitations in this study include the use of RNA expression data from published datasets without corresponding validation of protein expression. Nonetheless, the comprehensive nature of the analysis, which utilized all publicly available scRNA PCa datasets and the orthogonal validation in cell line and patient samples support the findings of uniform, high expression of B7H3 across PCa disease states as well as biochemical crosstalk between AR and B7H3 signaling. In addition, the role of AR in B7-H3 regulation has been explored only in a limited number of cell line models of PCa and needs further validation in preclinical models. Finally, miR29a is suspected to regulate B7-H3 expression[38] and could be another important regulator of B7-H3, confounding our understanding from current observations about AR and B7-H3, and requires further exploration.

## CONCLUSIONS

Our results establish B7-H3 as a suitable therapeutic target for not only primary prostatic adenocarcinoma, but also for castration resistant and NEPC. The broad and highly conserved expression of B7-H3 may possibly minimize tumor escape, via intratumoral heterogeneity, to B7-H3 targeted therapeutics. We believe our work (1) expands the present understanding about the influence of AR signaling cascade on B7-H3 expression and account for the effects of androgen ablation on the proliferative potential of tumors due to B7-H3 upregulation; (2) provides a framework to evaluate B7-H3 in multiple other cancer types; (3) improves inclusion/exclusion criteria for future B7-H3 directed PCa clinical trials.

## Authors’ Disclosures

E.S. is a paid consultant to Guidepoint Global, Gerson Lehrman Group (GLG), FirstThought, PathAI, CapitalOneSecurities, Boxer Capital, Health Advances, Forbion, Capvision, and Insighter. He receives institutional research funding from MacroGenics, Inc and AstraZeneca. The other authors have nothing to disclose.

## Authors’ Contributions

**S. Sharma**: Data curation, data visualization, formal analysis, validation, investigation, methodology, writing–original draft, writing–review and editing. **N. Mundhara:** Data curation, formal analysis, validation, investigation, methodology, writing–original draft, writing–review and editing. **E. Tekoglu:** Data curation, formal analysis, validation, investigation, methodology, writing–original draft, writing–review and editing. **A.A. Mendes:** Data curation, methodology, writing–review and editing. **T.L. Lotan**: Investigation, methodology, writing–review and editing. **P. Gu**: Data visualization. **J. Luo:** Investigation, methodology, writing–review and editing. **E.G. Baraban:** investigation, methodology, writing–review and editing. **N.A. Lack**: Data curation, formal analysis, supervision, funding acquisition, validation, investigation, methodology, writing–original draft, writing–review and editing. **E. Shenderov:** Conceptualization, resources, data curation, formal analysis, supervision, funding acquisition, validation, investigation, methodology, writing–original draft, writing–review and editing.

## Availability of Data

All bioinformatic data analysis is performed on publicly available data. We have developed an accessible graphical interface prostate single cell RNA data atlas visualizer available at Shenderovlab.com in the Tools Section. Code used for analysis is available at https://github.com/eshenderov/B7H3-PCa-Continuum

## Funding

This work was supported by TUBITAK 1001 (119Z279) to NAL and ET; Prostate Cancer Foundation Young Investigator Award (to E.S.); and Department of Defense grant (nos. W81XWH-16-PCRP-CCRSA and W81XWH-19-1-0511 to E.S.).

## Acknowledgements

We would like to thank Prof. Sushant Kachhap’s lab, Johns Hopkins School of Medicine for providing LNCaP and DU145 cells, Prof. Nate Brennen, Johns Hopkins School of Medicine for NCI-H660 and PrEC-LH cells, Dr. Rajendra Kumar’s lab, Johns Hopkins School of Medicine for VCaP, 22RV1, and PC3 cell lines, and protein lysates for LNCaP, VCaP, LAPC4, and 22RV1 control and R1881 (10 nM) treated for 96 h.

## REFERENCES

[1] Sung H, Ferlay J, Siegel RL, Laversanne M, Soerjomataram I, Jemal A, et al. Global Cancer Statistics 2020: GLOBOCAN Estimates of Incidence and Mortality Worldwide for 36 Cancers in 185 Countries. CA: A Cancer Journal for Clinicians. 2021;71:209–49.

[2] Siegel RL, Miller KD, Wagle NS, Jemal A. Cancer statistics, 2023. CA Cancer J Clin. 2023;73:17–48.

[3] Rebello RJ, Oing C, Knudsen KE, Loeb S, Johnson DC, Reiter RE, et al. Prostate cancer. Nature Reviews Disease Primers. 2021;7:9.

[4] Sartor O, Bono Jd, Chi KN, Fizazi K, Herrmann K, Rahbar K, et al. Lutetium-177–PSMA-617 for Metastatic Castration-Resistant Prostate Cancer. New England Journal of Medicine. 2021;385:1091–103.

[5] Kemble J, Kwon ED, Karnes RJ. Addressing the need for more therapeutic options in neuroendocrine prostate cancer. Expert Rev Anticancer Ther. 2023;23:177–85.

[6] Aggarwal V, Workman CJ, Vignali DAA. LAG-3 as the third checkpoint inhibitor. Nat Immunol. 2023;24:1415–22.

[7] Beer TM, Kwon ED, Drake CG, Fizazi K, Logothetis C, Gravis G, et al. Randomized, Double-Blind, Phase III Trial of Ipilimumab Versus Placebo in Asymptomatic or Minimally Symptomatic Patients With Metastatic Chemotherapy-Naive Castration-Resistant Prostate Cancer. J Clin Oncol. 2017;35:40–7.

[8] Powles T, Yuen KC, Gillessen S, Kadel EE, 3rd, Rathkopf D, Matsubara N, et al. Atezolizumab with enzalutamide versus enzalutamide alone in metastatic castration-resistant prostate cancer: a randomized phase 3 trial. Nat Med. 2022;28:144–53.

[9] Shenderov E, Boudadi K, Fu W, Wang H, Sullivan R, Jordan A, et al. Nivolumab plus ipilimumab, with or without enzalutamide, in AR-V7-expressing metastatic castration-resistant prostate cancer: A phase-2 nonrandomized clinical trial. Prostate. 2021;81:326–38.

[10] Sardinha M, Palma Dos Reis AF, Barreira JV, Fontes Sousa M, Pacey S, Luz R. Antibody-Drug Conjugates in Prostate Cancer: A Systematic Review. Cureus. 2023;15:e34490.

[11] Haffner MC, Zwart W, Roudier MP, True LD, Nelson WG, Epstein JI, et al. Genomic and phenotypic heterogeneity in prostate cancer. Nature reviews Urology. 2021;18:79–92.

[12] Vitale I, Shema E, Loi S, Galluzzi L. Intratumoral heterogeneity in cancer progression and response to immunotherapy. Nat Med. 2021;27:212–24.

[13] Yonesaka K, Haratani K, Takamura S, Sakai H, Kato R, Takegawa N, et al. B7-H3 Negatively Modulates CTL-Mediated Cancer Immunity. Clinical cancer research : an official journal of the American Association for Cancer Research. 2018;24:2653–64.

[14] Benzon B, Zhao SG, Haffner MC, Takhar M, Erho N, Yousefi K, et al. Correlation of B7-H3 with androgen receptor, immune pathways and poor outcome in prostate cancer: an expression-based analysis. Prostate Cancer Prostatic Dis. 2017;20:28–35.

[15] Shenderov E, De Marzo AM, Lotan TL, Wang H, Chan S, Lim SJ, et al. Neoadjuvant enoblituzumab in localized prostate cancer: a single-arm, phase 2 trial. Nature Medicine. 2023.

[16] Guo C, Figueiredo I, Gurel B, Neeb A, Seed G, Crespo M, et al. B7-H3 as a Therapeutic Target in Advanced Prostate Cancer. European urology. 2023;83:224–38.

[17] Mendes AA, Lu J, Kaur HB, Zheng SL, Xu J, Hicks J, et al. Association of B7-H3 expression with racial ancestry, immune cell density, and androgen receptor activation in prostate cancer. Cancer. 2022;128:2269–80.

[18] Brennen WN, Zhu Y, Coleman IM, Dalrymple SL, Antony L, Patel RA, et al. Resistance to androgen receptor signaling inhibition does not necessitate development of neuroendocrine prostate cancer. JCI Insight. 2021;6.

[19] Ghandi M, Huang FW, Jané-Valbuena J, Kryukov GV, Lo CC, McDonald ER, et al. Next-generation characterization of the Cancer Cell Line Encyclopedia. Nature. 2019;569:503–8.

[20] Hieronymus H, Lamb J, Ross KN, Peng XP, Clement C, Rodina A, et al. Gene expression signature-based chemical genomic prediction identifies a novel class of HSP90 pathway modulators. Cancer cell. 2006;10:321–30.

[21] Beltran H, Prandi D, Mosquera JM, Benelli M, Puca L, Cyrta J, et al. Divergent clonal evolution of castration-resistant neuroendocrine prostate cancer. Nat Med. 2016;22:298–305.

[22] Hänzelmann S, Castelo R, Guinney J. GSVA: gene set variation analysis for microarray and RNA-Seq data. BMC Bioinformatics. 2013;14:7.

[23] Dobin A, Davis CA, Schlesinger F, Drenkow J, Zaleski C, Jha S, et al. STAR: ultrafast universal RNA-seq aligner. Bioinformatics. 2012;29:15–21.

[24] Love MI, Huber W, Anders S. Moderated estimation of fold change and dispersion for RNA-seq data with DESeq2. Genome biology. 2014;15:550.

[25] Hao Y, Stuart T, Kowalski MH, Choudhary S, Hoffman P, Hartman A, et al. Dictionary learning for integrative, multimodal and scalable single-cell analysis. Nature biotechnology. 2024;42:293–304.

[26] McGinnis CS, Murrow LM, Gartner ZJ. DoubletFinder: Doublet Detection in Single-Cell RNA Sequencing Data Using Artificial Nearest Neighbors. Cell Systems. 2019;8:329–37.e4.

[27] Lotan TL, Gupta NS, Wang W, Toubaji A, Haffner MC, Chaux A, et al. ERG gene rearrangements are common in prostatic small cell carcinomas. Mod Pathol. 2011;24:820–8.

[28] Tan HL, Sood A, Rahimi HA, Wang W, Gupta N, Hicks J, et al. Rb loss is characteristic of prostatic small cell neuroendocrine carcinoma. Clinical cancer research : an official journal of the American Association for Cancer Research. 2014;20:890–903.

[29] Culig Z, Santer FR. Androgen receptor signaling in prostate cancer. Cancer Metastasis Rev. 2014;33:413–27.

[30] Cai C, He Housheng H, Chen S, Coleman I, Wang H, Fang Z, et al. Androgen Receptor Gene Expression in Prostate Cancer Is Directly Suppressed by the Androgen Receptor Through Recruitment of Lysine-Specific Demethylase 1. Cancer cell. 2011;20:457–71.

[31] Verras M, Lee J, Xue H, Li T-H, Wang Y, Sun Z. The Androgen Receptor Negatively Regulates the Expression of c-Met: Implications for a Novel Mechanism of Prostate Cancer Progression. Cancer Research. 2007;67:967–75.

[32] Bhatia V, Kamat NV, Pariva TE, Wu L-T, Tsao A, Sasaki K, et al. Targeting advanced prostate cancer with STEAP1 chimeric antigen receptor T cell and tumor-localized IL-12 immunotherapy. Nature Communications. 2023;14:2041.

[33] Ajkunic A, Sayar E, Roudier MP, Patel RA, Coleman IM, De Sarkar N, et al. Assessment of TROP2, CEACAM5 and DLL3 in metastatic prostate cancer: Expression landscape and molecular correlates. npj Precision Oncology. 2024;8:104.

[34] Brady L, Kriner M, Coleman I, Morrissey C, Roudier M, True LD, et al. Inter- and intra-tumor heterogeneity of metastatic prostate cancer determined by digital spatial gene expression profiling. Nature Communications. 2021;12:1426.

[35] Sayar E, Patel RA, Coleman IM, Roudier MP, Zhang A, Mustafi P, et al. Reversible epigenetic alterations mediate PSMA expression heterogeneity in advanced metastatic prostate cancer. JCI Insight. 2023;8.

[36] Nolan-Stevaux O, Li C, Liang L, Zhan J, Estrada J, Osgood T, et al. AMG 509 (Xaluritamig), an Anti-STEAP1 XmAb 2+1 T-cell Redirecting Immune Therapy with Avidity-Dependent Activity against Prostate Cancer. Cancer Discovery. 2024;14:90–103.

[37] Aurilio G, Cimadamore A, Mazzucchelli R, Lopez-Beltran A, Verri E, Scarpelli M, et al. Androgen Receptor Signaling Pathway in Prostate Cancer: From Genetics to Clinical Applications. Cells. 2020;9.

[38] Xu H, Cheung IY, Guo HF, Cheung NK. MicroRNA miR-29 modulates expression of immunoinhibitory molecule B7-H3: potential implications for immune based therapy of human solid tumors. Cancer Res. 2009;69:6275–81.

