## Supplementary for "B7-H3 as a Dual Clinically Relevant Checkpoint and Antibody Drug Conjugate Target Expressed Across Adenocarcinoma and Neuroendocrine Prostate Cancers"

### Supplementary Tables

| Reagent | Company | CatLog No. |
|---|---|---|
| Phenol free RPMI media | Gibco | 11835030 |
| Heat inactivated Fetal bovine Serum | Gibco | 16140071 |
| Charcoal stripped FBS | Gibco | 12676029 |
| 10X Cell lysis buffer | Cell Signaling Technology | 9803S |
| Protease Inhibitor cocktail | Roche | 1183615301 |
| Phosphatase Inhibitor | Roche | 4906845001 |
| Anti-B7-H3 antibody (Clone D9M2L) | Cell Signaling Technology | CST#14058 |
| Anti-PD-L1 antibody (Clone E1L3N) | Cell Signaling Technology | CST#13684 |
| Anti-PD-L2 antibody (Clone 176611) | ThermoScientific | MA5-23887 |
| Anti-STEAP1 antibody (Clone B-4) | Santa Cruz Biotechnology | sc-271872 |
| Anti-STEAP2 polyclonal antibody | ThermoScientific | 20201-1-AP |
| Anti-DLL3 antibody (Clone 11H27L9) | ThermoScientific | 703075 |
| Anti-Nectin1 antibody (Clone CK8) | Invitrogen | 10168972 |
| Anti-PSCA polyclonal antibody | Invitrogen | PIPA575178 |
| Anti-41BB antibody (Clone – EPR20238) | Abcam | ab209256 |
| Anti-TROP2 antibody (Clone – EPR20043) | Abcam | ab214488 |
| Anti-LAG3 antibody (Clone – EPR20261) | Abcam | ab209236 |
| Anti-PSMA antibody (Clone GCP-04) | NOVUS Biologicals | NBP1-45057 |
| Anti-β-actin antibody | Abcam | ab8226 |
| Anti-AR antibody | Santa Cruz Biotechnology | sc-7305 |
| R1881 | PerkinElmer | NLP005005MG |
| Enzalutamide | Medchem Express | HY-70002 |
| Qiagen RNeasy kit | Qiagen | 74004 |
| High-Capacity cDNA Reverse Transcription Kit with RNase Inhibitor | ThermoScientific | Cat# 4374966 |
| TaqMan probe c-MET | ThermoScientific | Hs00179845_m1 |
| TaqMan probe CD276 | ThermoScientific | Hs00987207_m1 |
| TaqMan probe AR | ThermoScientific | Hs00171172_m1 |
| TaqMan probe KLK2 | ThermoScientific | Hs00428384_g1 |
| TaqMan probe GAPDH | ThermoScientific | Hs02786624_g1 |
| Fc receptor binding inhibitor | eBioscience | 14-9161-73 |
| LIVE/DEAD™ Fixable Aqua Dead Cell stain kit | ThermoScientific | 34966) |
| Anti-AR AF488 antibody | Cell Signaling Technology | ST#7395 |
| FOXP3 transcription factor staining kit | Invitrogen | 00-5521-00 |

**Supplementary Table S1:** Reagents, antibodies, TaqMan probes vendor and catalog numbers

| | Reference number | Condition |
|---|---|---|
| 1 | GSE168671 | LAPC4 1nM R1881 4h, LNCaP 1nM R1881 4h |
| 2 | GSE148397 | LAPC4 1nM R1881 8h, LAPC4 1nM R1881 22h, VCaP 1nM R1881 8h, VCaP 1nM R1881 22h |
| 3 | GSE13627 | LNCaP 1nM R1881 12h, VCaP 1nM R1881 12h |
| 4 | GSE137056 | LNCaP 1nM R1881 24h |
| 5 | GSE159606 | LNCaP 10nM DHT 16h |
| 6 | GSE161170 | LNCaP 10nM R1881 16h |
| 7 | GSE163539 | LNCaP 10nM DHT 24h |
| 8 | GSE223024 | VCaP 10nM DHT 24h |
| 9 | GSE64529 | LNCaP 100nM DHT 6h |
| 10 | GSE83652 | VCaP 1nM DHT 16h |

**Supplementary Table S2:** Publicly available RNA-seq datasets obtained from Gene Expression Omnibus (GEO) for multiple cell lines and treatment conditions

| | Dataset | Subtype | # Patients | Cell Count | UMI (Mean) | Genes (Mean) | Mito RNA (Mean) |
|---|---|---|---|---|---|---|---|
| 1 | GSE120716 | Healthy | 3 | 26893 | 4774 | 1430 | 3.95 |
| 2 | GSE137829 | CRPC | 3 | 6907 | 16270 | 3046 | 4.65 |
| 3 | GSE137829 | NEPC | 2 | 8456 | 9889 | 2034 | 4.13 |
| 4 | GSE137829 | Small Cell | 1 | 8179 | 13361 | 2679 | 5.91 |
| 5 | GSE141445 | Adeno | 6 | 15750 | 10586 | 2348 | 6.57 |
| 6 | GSE141445 | ICC/IDC | 7 | 40103 | 8390 | 1793 | 11.64 |
| 7 | GSE176031 | Adeno | 6 | 10755 | 3984 | 1732 | 7.26 |
| 8 | GSE176031 | Benign | 4 | 4223 | 2983 | 1435 | 5.68 |
| 9 | GSE176031 | ICC/IDC | 5 | 5177 | 3387 | 1615 | 5.22 |
| 10 | GSE181294 | Adeno | 18 | 62989 | 3394 | 1137 | 0.00 |
| 11 | GSE181294 | Benign | 14 | 61482 | 2775 | 957 | 0.00 |
| 12 | GSE181294 | Healthy | 5 | 17783 | 2728 | 860 | 0.00 |
| 13 | GSE185344 | ICC/IDC | 7 | 55310 | 8845 | 2164 | 4.68 |
| 14 | GSE193337 | Benign | 4 | 11019 | 6887 | 1897 | 10.31 |
| 15 | GSE193337 | ICC/IDC | 4 | 12188 | 10457 | 2407 | 11.18 |
| 16 | GSE244823 | Adeno | 10 | 98850 | 6393 | 1888 | 4.20 |
| 17 | HRA002145 | Adeno | 1 | 4720 | 10116 | 2352 | 6.47 |
| 18 | HRA002145 | NEPC | 2 | 7605 | 6222 | 1767 | 4.72 |
| 19 | HRA002145 | Small Cell | 1 | 4428 | 8825 | 2607 | 9.01 |
| 20 | GSE264573 | Adeno | 8 | 41489 | 8325 | 2954 | 0.00 |
| 21 | GSE264573 | CRPC | 10 | 46107 | 8538 | 2976 | 0.00 |
| 22 | GSE264573 | Small Cell | 3 | 13856 | 10308 | 3829 | 0.00 |

**Supplementary Table S3**: Characterization of the single cell RNA sequencing PCa datasets obtained after performing quality control measures. Eleven single-cell datasets encompassing Healthy, Adjacent Benign, BPH, Primary PCa, CRPC and NEPC PCa subtypes. All the datasets excluding GSE120716 (Healthy and GSE141445 (ICC/IDC) are composed of multiple subtypes of Prostate Cancer.

| | Cell Type | Markers |
|---|---|---|
| 1 | Epithelial | KRT8, KRT18 |
| 2 | Mesenchymal | IGFBP7, SPARCL1 |
| 3 | Immune | RGS1, PTPRC, LAPTM5, ALOX5AP, GPR183, SAMSN1 |
| 4 | Neuronal | PLP1, MPZ |
| 5 | Luminal | KLK2, KLK3, TMPRSS2, FOXA1, SLC45A3, RDH11, NKX3-1 |
| 6 | Basal | KRT5, KRT14, KRT15, KRT17, DST |
| 7 | Club | SCGB3A1, LCN2, PIGR, WFDC2, MMP7 |
| 8 | Hillock | KRT13, GPX2, S100P |
| 9 | Neuroendocrine | CHGA, CHGB, CALCA, SCG2, ENO2, LMO3, SCGN, TMEM61, ASCL1, PEG10, SEC11C |
| 10 | Fibroblast | FBLN1, CFD, SERPINF1, PTN, MMP2, DCN, LUM, COL1A1, COL1A2, IGF1, C7, CCDC80, LTBP4, PDGFRA, FBLN2 |
| 11 | Pericyte | THY1, ANGPT2, COL3A1, COL5A2 |
| 12 | Endothelial | CLDN5, IFI27, VWF, PECAM1, FLT1, PTPRB, PLVAP, ENG, RAMP2, AQP1 |
| 13 | Smooth | Muscle, ACTA2, RGS5, MYH11, TAGLN, MYL9, MYLK, MCAM |
| 14 | Myeloid | LYZ, BCL2A1, AIF1, MS46A, FCGR2A, MS4A7, LST1, IL1B, IFI30, CD68, CD1C, FCER1A, S100A8, S100A9, FCGR3A, CD14 |
| 15 | T | CD3E, CD3D, CD3G, IL7R, CD2, CD7 |
| 16 | NK | NKG7, GZMB, KLRD1 |
| 17 | B | MS4A1, CXCR5, BANK1, LY9 |
| 18 | Plasma | IGKC, IGHA1, IGJ, IGHA2, AC096579.7, MZB1, IGHG3, IGHG4, IGHG1, JCHAIN |
| 19 | Mast | KIT, CPA3, TPSAB1, VWA5A, IL1RL1, SLC18A2, MS4A2, TPSB2 |

**Supplementary Table S4**: Characteristic markers of cell types annotated in the integrated single cell RNA sequencing dataset. Epithelial, Mesenchymal, Immune and Neuronal are the lineages and the remaining are the subtypes that the cells are annotated with. Neuronal cells were explored using the mentioned markers, but were not observed. These markers are an ensemble of the markers reported in the PCa scRNA-seq datasets in this work.

|  | Targets | Bulk RNA sequencing | Single cell RNA sequencing |
|---|---|---|---|
| 1 | B7-H3 | 12.10% | 6.23% |
| 2 | NECTIN1 | 26.70% | 141.85% |
| 3 | FKBP5 | 31.10% | 9.03% |
| 4 | IGF1R | 32.90% | 6.78% |
| 5 | STEAP1 | 43.20% | 14.39% |
| 6 | TROP2 | 44.70% | 11.08% |
| 7 | LAG3 | 69.90% | 21.26% |
| 8 | STEAP2 | 70.20% | 15.48% |
| 9 | PSCA | 80.40% | 31.76% |
| 10 | PD-L1 | 85.50% | 71.12% |
| 11 | TMPRSS2 | 87.40% | 12.70% |
| 12 | DLL3 | 94.10% | 38.23% |
| 13 | PD-L2 | 127% | 334.52% |
| 14 | OX40 | 138% | 46.42% |
| 15 | PSMA | 146% | 14.30% |
| 16 | AR | 150% | 10.86% |
| 17 | KLK2 | 150% | 18.44% |
| 18 | KLK3 | 157% | 21.11% |
| 19 | TIGIT | 173% | 56.47% |
| 20 | 4-1BB | 198% | 64.66% |
| 21 | PD-1 | 230% | 37.39% |
| 22 | CTLA4 | 269% | 54.93% |

**Supplementary Table S5:** Coefficient of variation for ADC and ICI investigated in this study in bulk and single cell RNA sequencing datasets. Among the tumor-specific markers, and consistent with analyzed cell lines, B7-H3 consistently showed the least amount of variation in CCLE bulk RNA sequencing dataset (12.10%) and single-cell RNA sequencing integrated dataset (CV = 6.23%). This was followed by NECTIN1 in bulk RNA sequencing data (CV = 26.70%) and IGF1R in single-cell RNA sequencing data (CV = 6.77%). Other tumor-specific markers showed higher variability. Sorted by bulk RNAseq data.

### Supplementary Figures

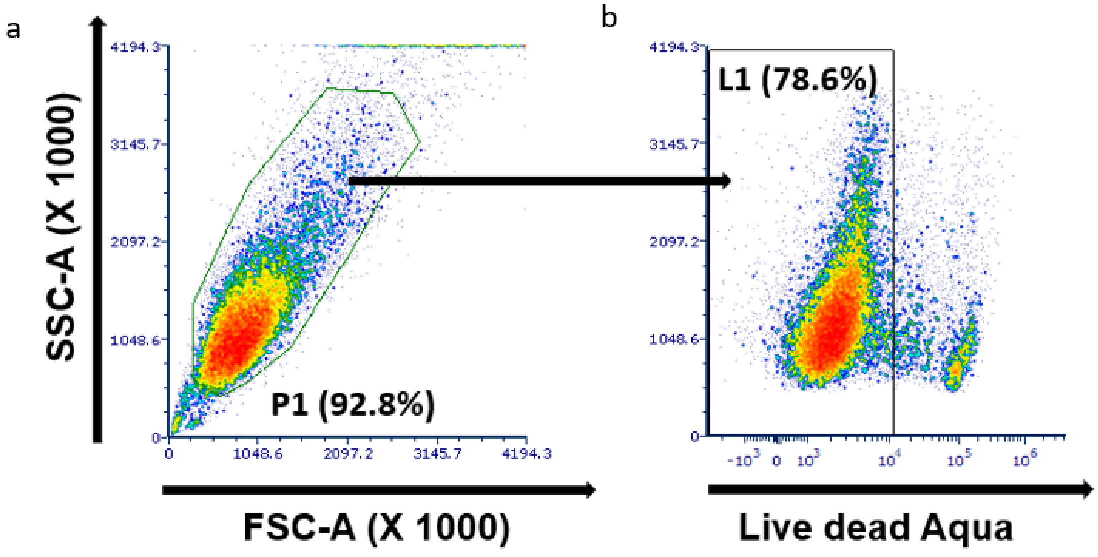


**Supplementary Figure S1: Gating strategy for flow cytometry analysis using FCS express**. (a) SSC VS FSC plot was gated for P1 population to remove the necrotic cells. (b) The P1 population was gated on live cells (L1) using live dead aqua dye. These selected L1 population was used for further analysis of AR, B7-H3 expression levels in LNCaP, DU145 and 22RV1 cells as a part of Fig (4b & 4e) of the manuscript.

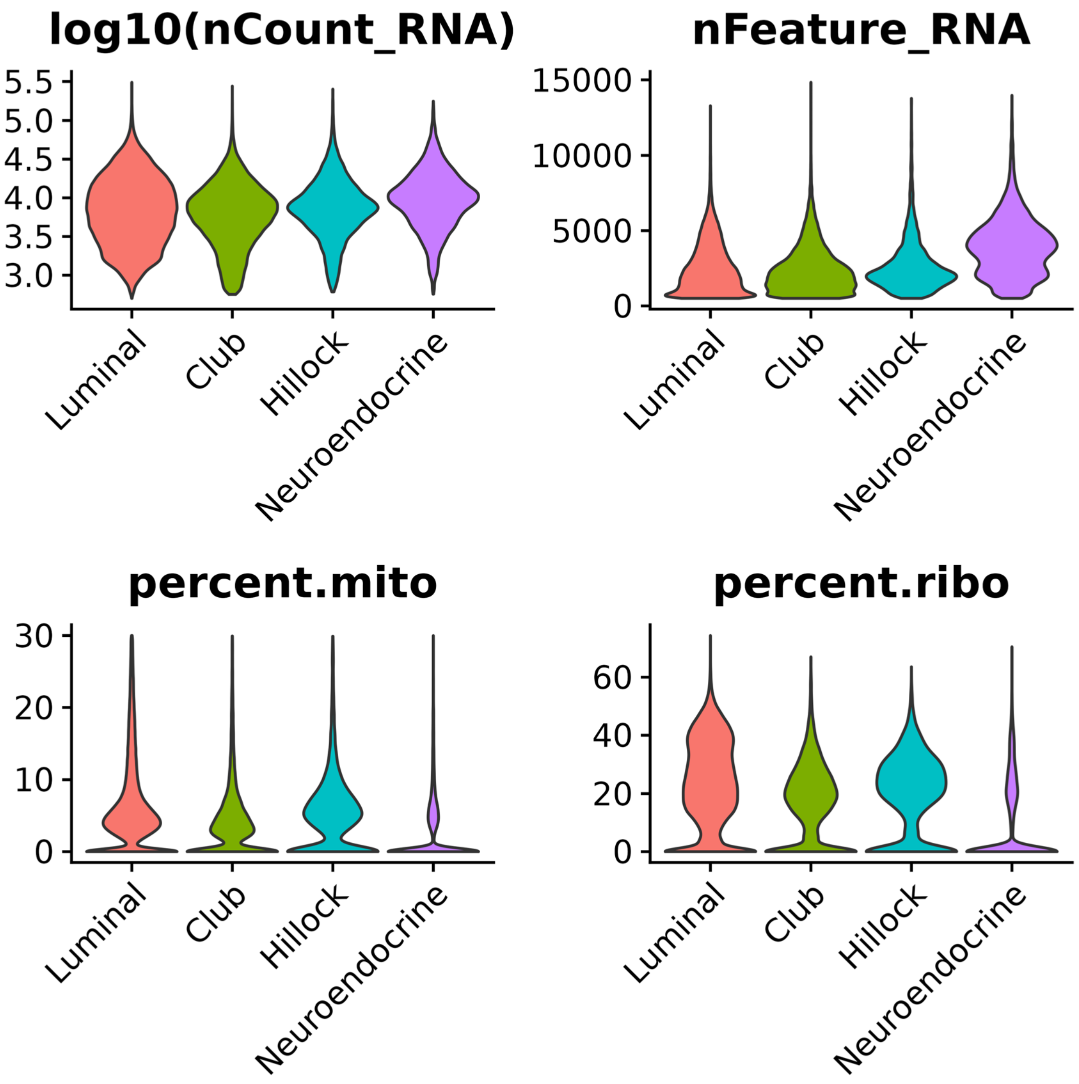


**Supplementary Figure S2**: Integrated single cell prostate dataset was filtered for cells with number of genes (nFeature_RNA) > 500 and percentage of mitochondrial RNA (percent.mt) < 25%. The distribution of these two metrics, along with log10 of total RNA counts (nCounts_RNA) and percentage of ribosomal RNA, is shown in the figure for Luminal, Club, Hillock and Neuroendocrine cells. No cutoffs were applied to total RNA counts or ribosomal percentage, but displayed here for

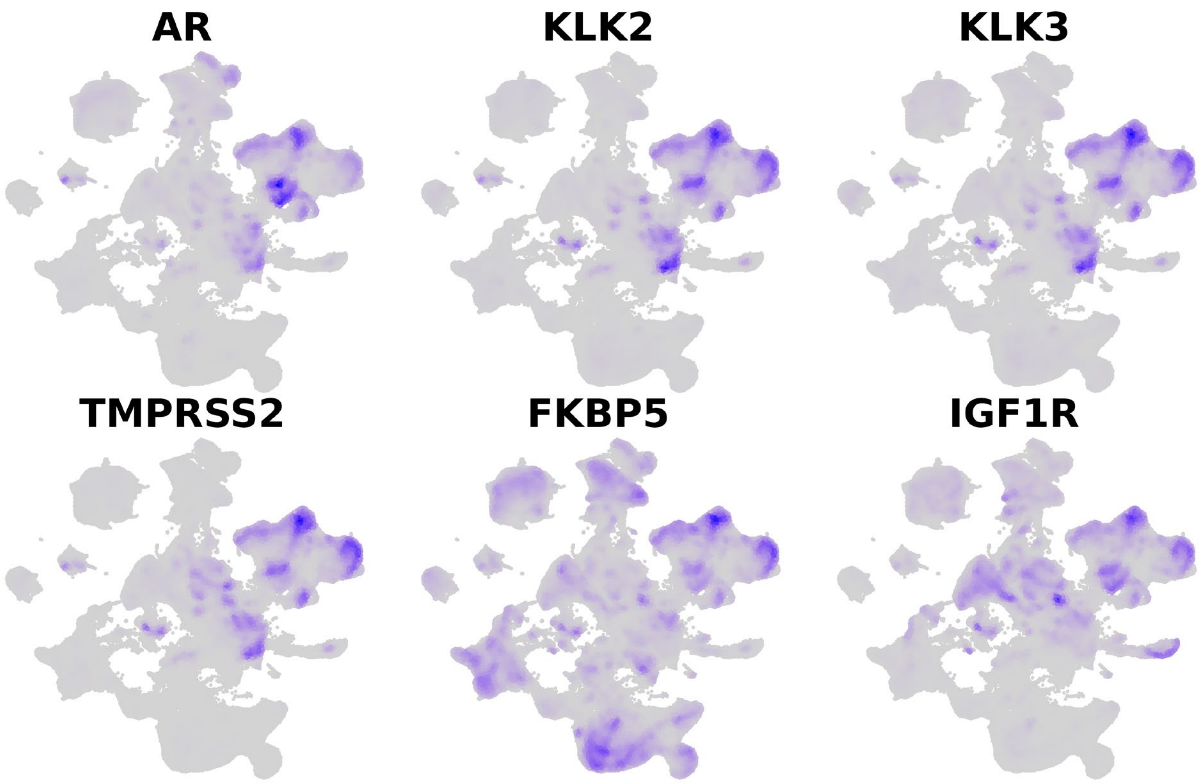


**Supplementary Figure S3**: Expression densities of prostate epithelial markers in the integrated single-cell PCa dataset. The markers are negative for basal cells, which is not an observed cell type in tumor regions.

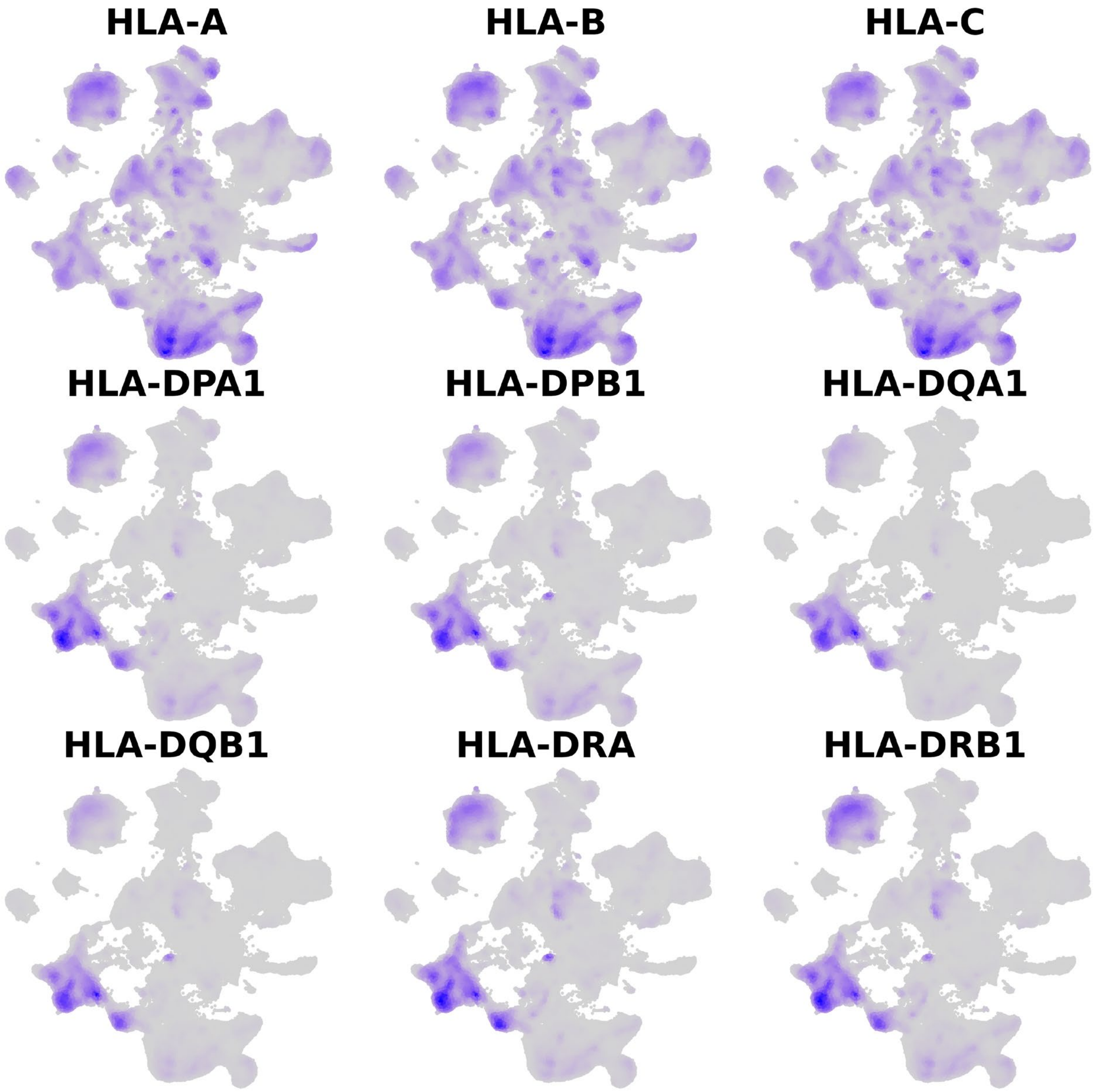


**Supplementary Figure S4**: Expression densities of HLA genes in the integrated single-cell PCa dataset

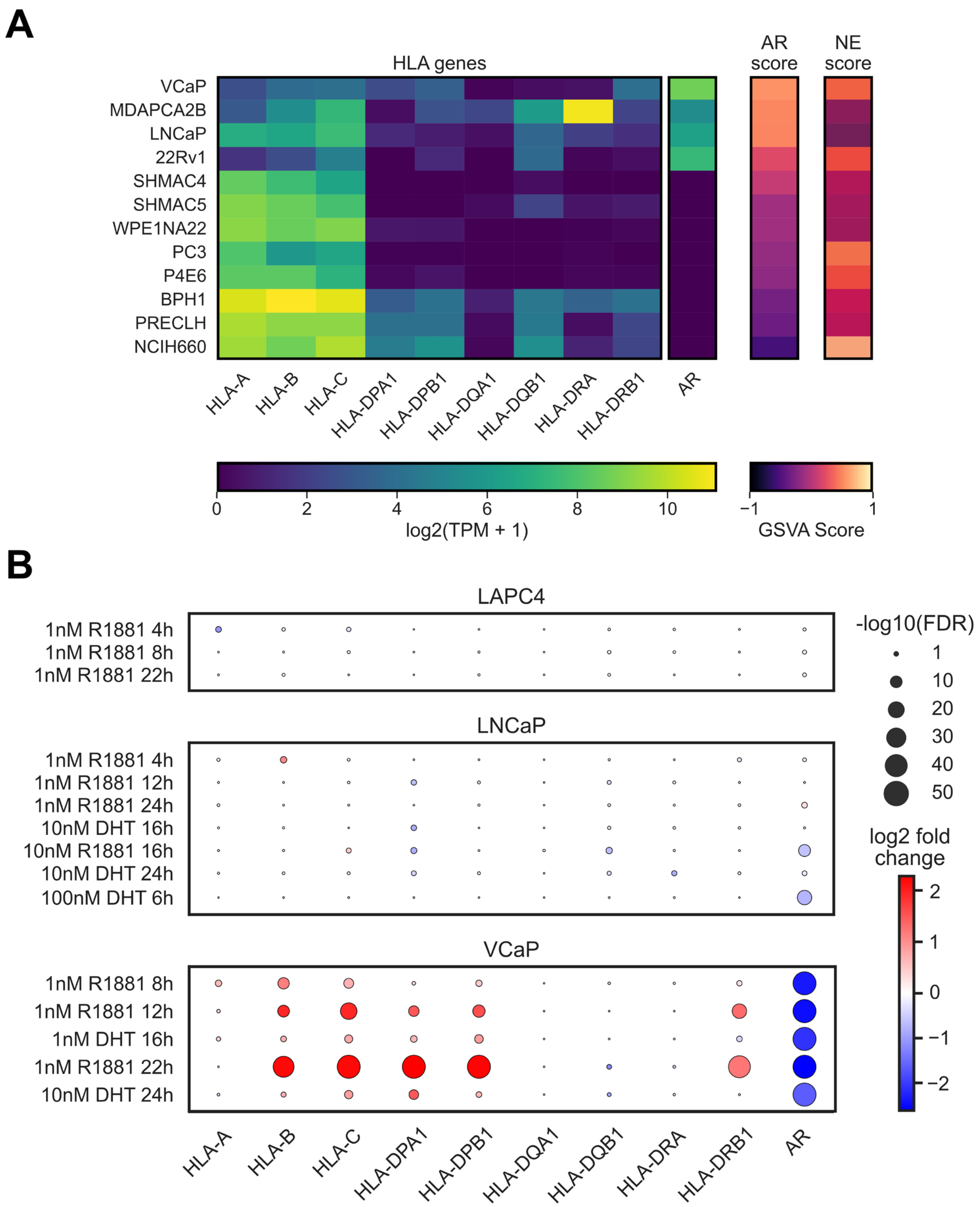


**Supplementary Figure S5: (A)** Log TPM expression of HLAs in prostate cancer cell lines, with their corresponding AR and NE scores. (**B**) Differential gene expression of HLA with androgen treatment in LAPC4, LNCaP and VCAP cell lines. Fold-change in HLAs is shown upon treatment with different concentrations and treatment durations of R1881 and DHT.

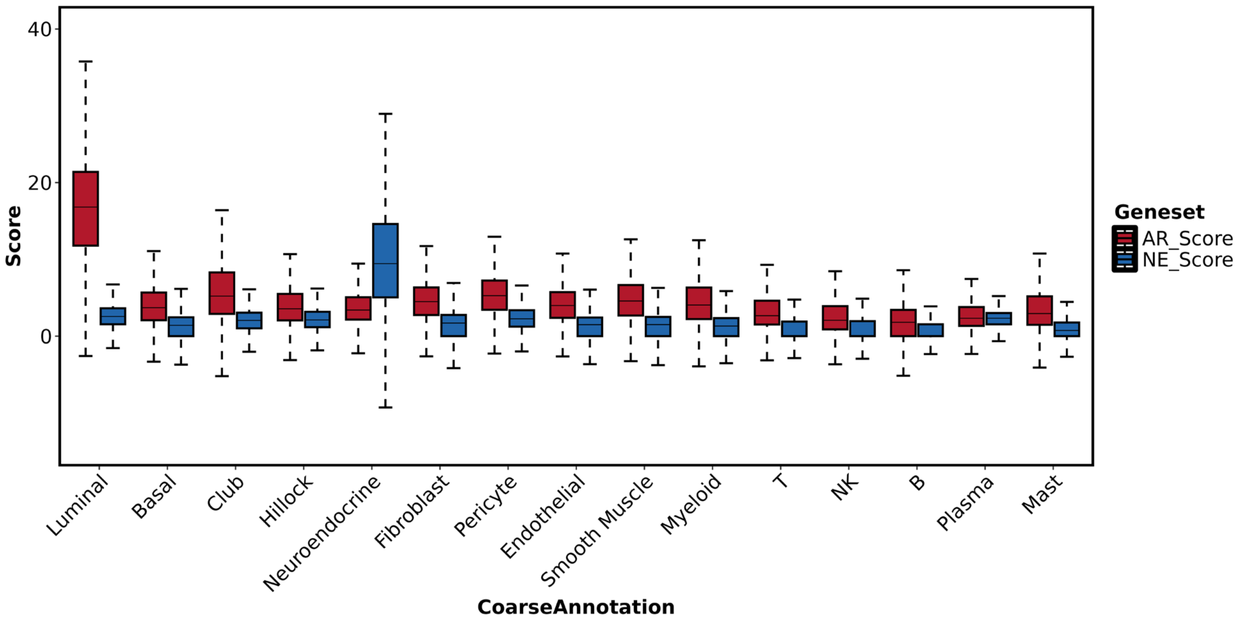


**Supplementary Figure S6**: AR and NE score for cellular subtypes in the integrated single-cell PCa dataset, indicating highest AR and NE score for Luminal and Neuroendocrine clusters of the UMAP of the integrated PCa scRNA-seq dataset.

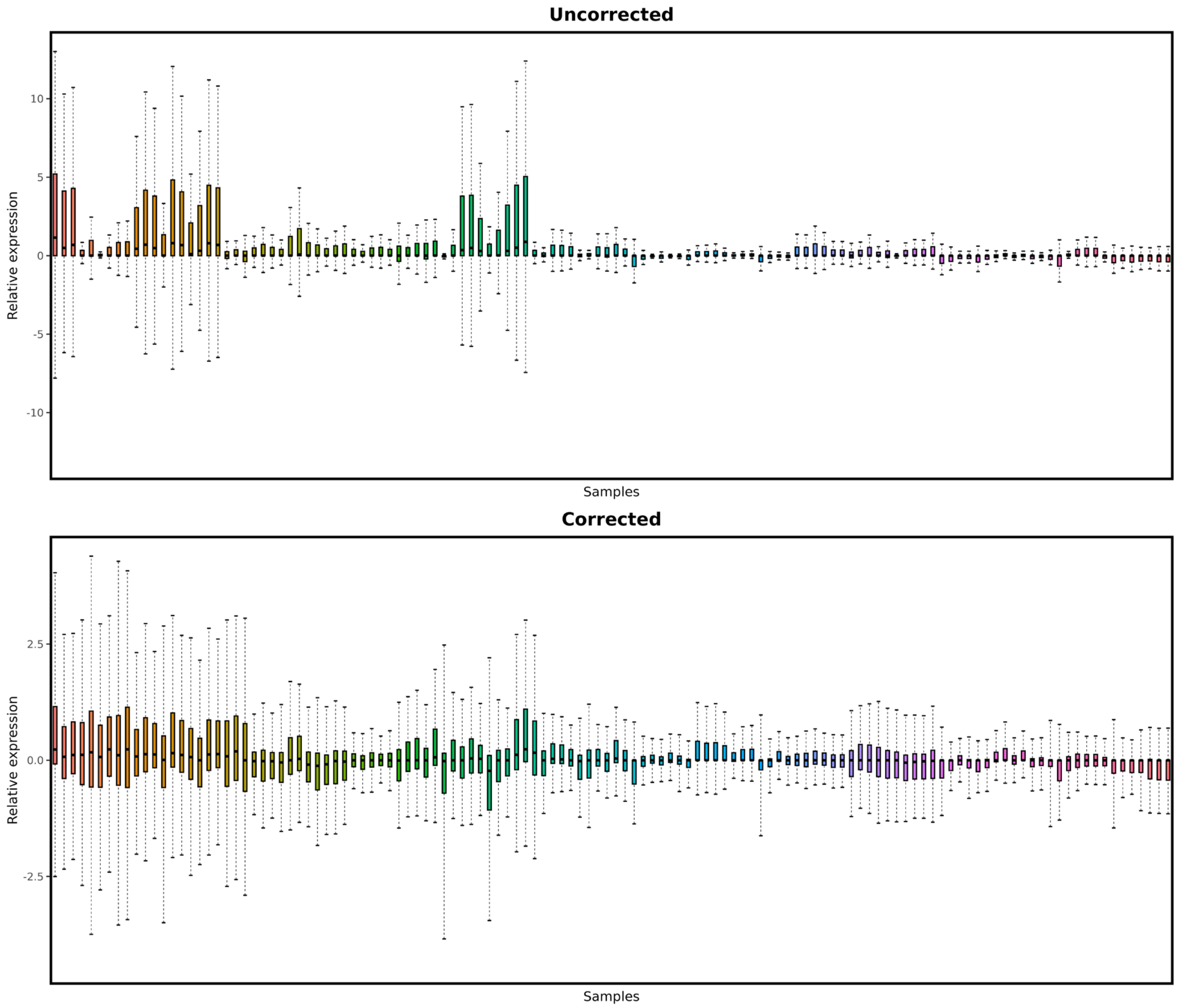


**Supplementary Figure S7**: Integrated single cell prostate dataset was subsetted for clusters with highest AR and NE score. The subsetted dataset was converted into pseudobulk dataset and the resulting pseudobulk dataset was batch-corrected using limma. **A**) Uncorrected pseudobulk log counts, **B**) Batch-corrected pseudobulk log counts.

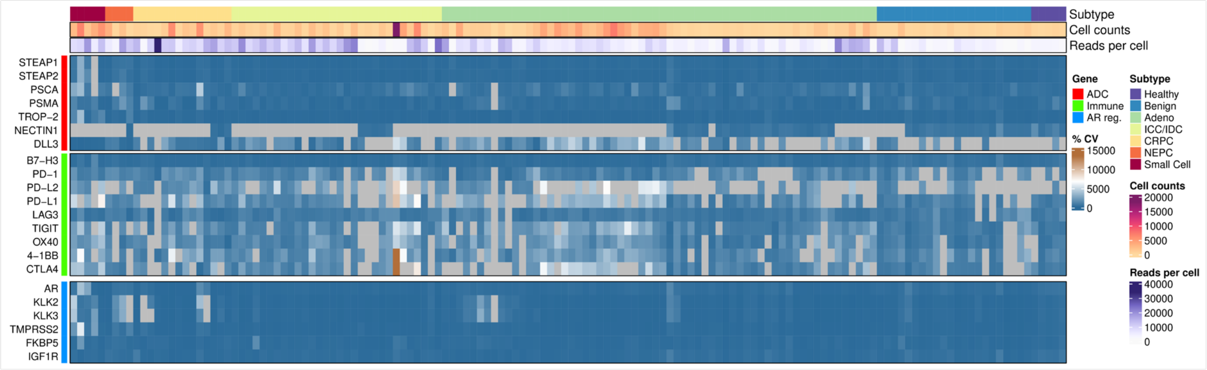


**Supplementary Figure S8**: Percent coefficient of variation (% CV) of ADC targets across Luminal and Neuroendocrine cell populations in each patient. %CV is defined as the ratio of standard deviation to mean of the normalized expression of a target in a patient sample. %CV is jointly calculated over all the cells clustered in Luminal and Neuroendocrine.

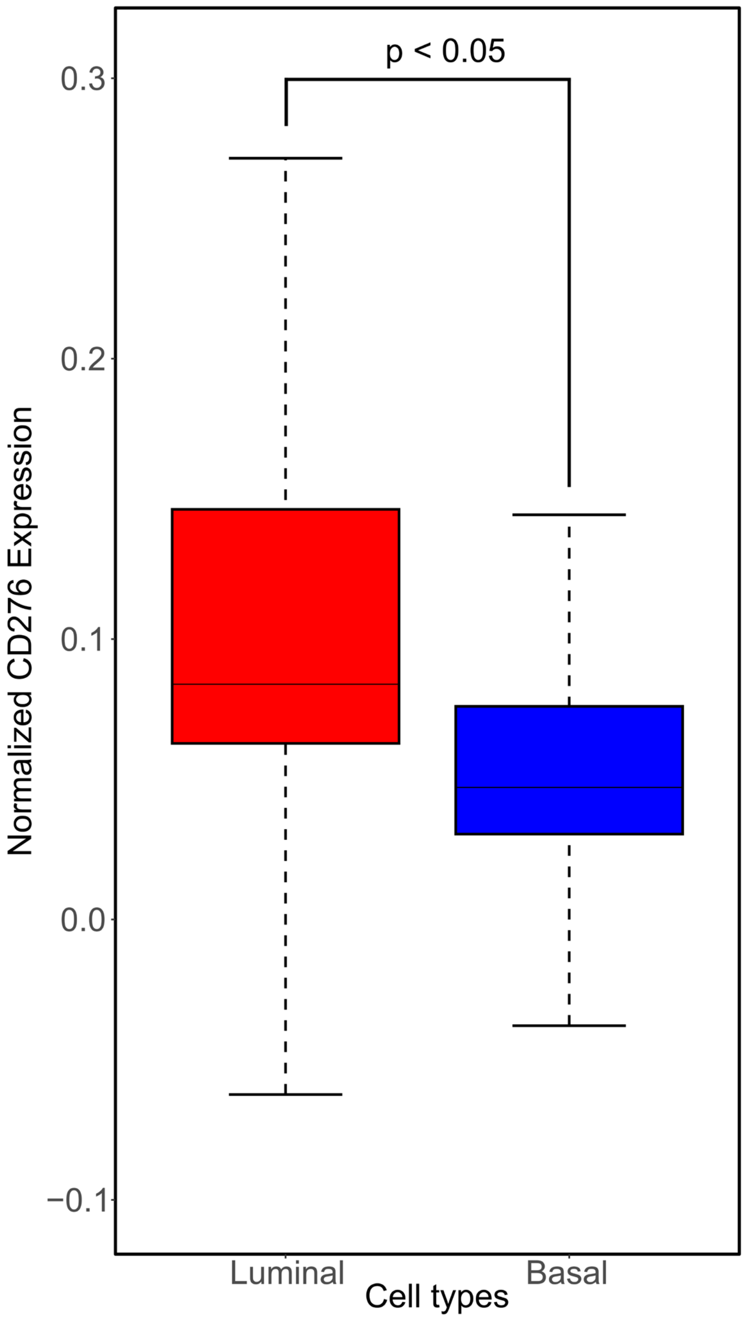


**Supplementary Figure S9**: Luminal cell have higher RNA expression of B7-H3 (CD276) than basal cells in the healthy and benign samples of integrated single cell prostate dataset. Exact p-value is 0.0115 and was obtained from Welch t-test in R.

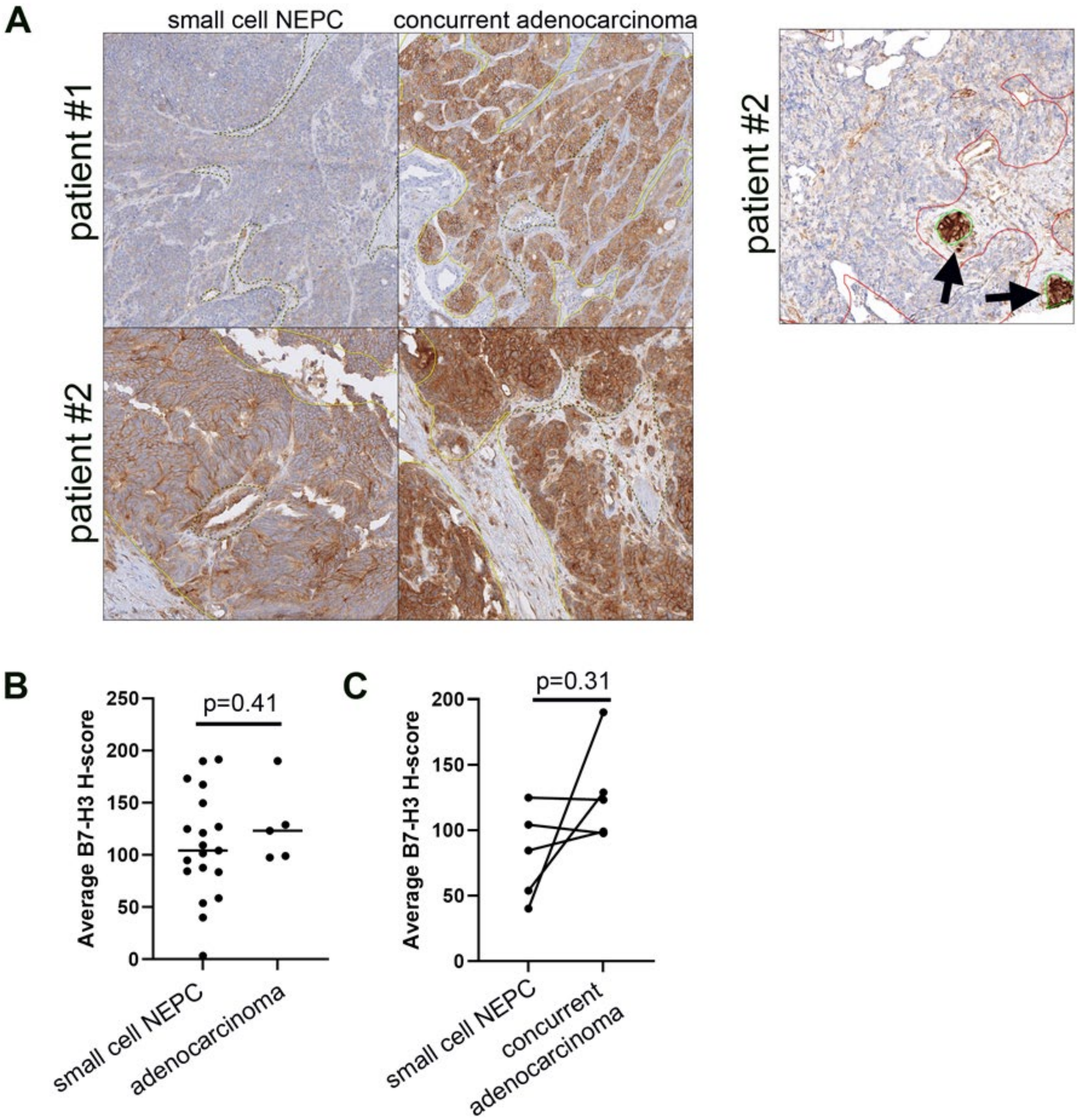


**Supplementary Figure S10**: (**A**) Representative patients with NEPC and concurrent prostatic acinar adenocarcinoma from a cohort of 29 prostatic small cell carcinoma specimen. (**B**) Comparison of average B7-H3 H-score between neuroendocrine (NEPC) and primary PCa specimen. (**C**) Paired comparison of average B7-H3 H-score between neuroendocrine and concomitant acinar adenocarcinoma components across 5 patients.

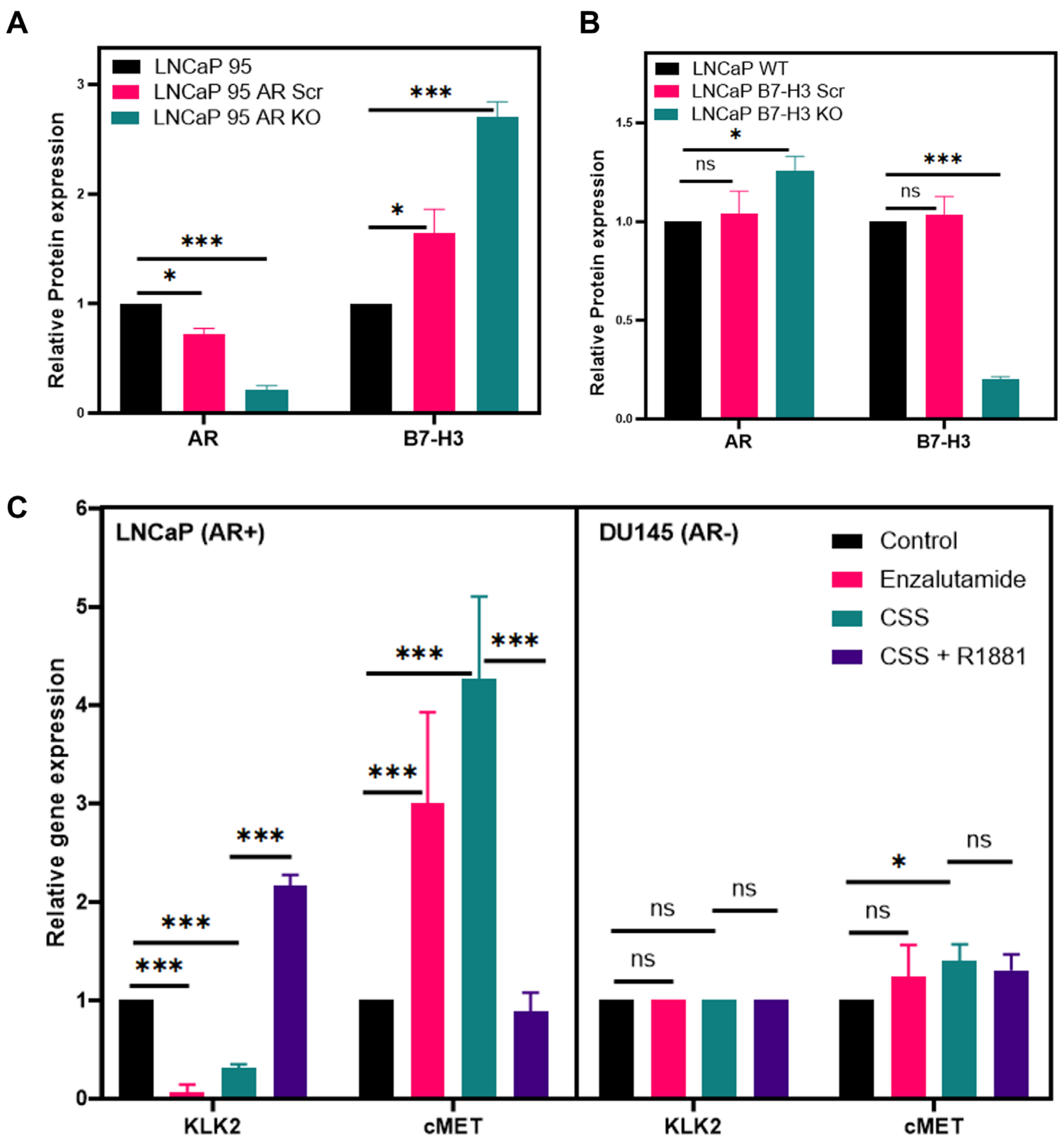


**Figure S11:** (**A**) Densitometric quantification of the western blot shown in figure 3c. The data is represented with respect to the LNCaP 95 cell line. All data points are normalized to actin (**B**) Densitometric quantification of the western blot shown in Figure 3c. The data is represented with respect to the LNCaP WT cell line. All data points are normalized to action. (**C**) Relative gene expression of KLK2, c-MET in LNCaP and DU145 cells. Bar graph representing the relative gene expression upon treatment with Enzalutamide (10 uM), CSS, CSS + R1881 (10 nM) for 96 h (N=3) with respect to RPMI untreated controls. All the data points are normalized to GAPDH control. The error bars represent SD (ns = non-significant, *** = p value < 0.001, ** = p value < 0.01, * = p value < 0.05).
